# Development of a small molecule that corrects misfolding and increases secretion of Z α_1_-antitrypsin

**DOI:** 10.1101/2020.07.26.217661

**Authors:** David A. Lomas, James A. Irving, Christopher Arico-Muendel, Svetlana Belyanskaya, Andrew Brewster, Murray Brown, Chun-wa Chung, Hitesh Dave, Alexis Denis, Nerina Dodic, Anthony Dossang, Peter Eddershaw, Diana Klimaszewska, Imran Haq, Duncan S. Holmes, Jonathan P. Hutchinson, Alistair Jagger, Toral Jakhria, Emilie Jigorel, John Liddle, Ken Lind, Stefan J. Marciniak, Jeff Messer, Margaret Neu, Allison Olszewski, Adriana Ordonez, Riccardo Ronzoni, James Rowedder, Martin Rüdiger, Steve Skinner, Kathrine J. Smith, Rebecca Terry, Lionel Trottet, Iain Uings, Steve Wilson, Zhengrong Zhu, Andrew C. Pearce

## Abstract

Severe α_1_-antitrypsin deficiency results from the Z allele (Glu342Lys) that causes the accumulation of homopolymers of mutant α_1_-antitrypsin within the endoplasmic reticulum of hepatocytes in association with liver disease. We have used a DNA-encoded chemical library to undertake a high throughput screen to identify small molecules that bind to, and stabilise Z α_1_-antitrypsin. The lead compound blocks Z α_1_-antitrypsin polymerisation *in vitro*, reduces intracellular polymerisation and increases the secretion of Z α_1_-antitrypsin three-fold in mammalian cells including an iPSC model of disease. Crystallographic and biophysical analyses demonstrate that GSK716 and related molecules bind to a cryptic binding pocket, negate the local effects of the Z mutation and stabilise the bound state against progression along the polymerization pathway. Oral dosing of transgenic mice at 100 mg/kg three times a day for 20 days increased the secretion of Z α_1_-antitrypsin into the plasma by 7-fold. There was no observable clearance of hepatic inclusions with respect to controls. This study provides proof-of-principle that ‘mutation ameliorating’ small molecules are a viable approach to treat protein conformational diseases.

## Introduction

Alpha-1 antitrypsin deficiency affects 1 in 2000 people of Northern European descent, leading to liver and lung disease (*1*). Ninety-five percent of severe deficiency results from the ‘Z’ allele (Glu342Lys) that perturbs the folding of α_1_-antitrypsin resulting in the secretion of only 15% of the mature protein. The remaining protein is retained within the cell by persistent binding to molecular chaperones (*2*) and then either degraded via the ERAD-proteasome pathway (*3–5*) or folded into ordered polymers that may be cleared by autophagy (*6*) or accumulate within the endoplasmic reticulum (ER) of hepatocytes (*7*). The accumulation of polymers causes neonatal hepatitis, cirrhosis and hepatocellular carcinoma, and can sensitise the liver to damage from environmental insults such as alcohol, fat or viral hepatitis (*8, 9*). The consequent deficiency of α_1_-antitrypsin within the circulation results in insufficient protection of the lungs from neutrophil elastase, leading to early onset emphysema (*1*).

The Z mutation lies at the head of strand 5 of β-sheet A of α_1_-antitrypsin. It perturbs the local environment, allowing population of an unstable intermediate that we have termed M* (*10*) in which β-sheet A opens and the upper part of helix F unwinds (*11, 12*). Polymerisation from this state involves insertion of the RCL into β-sheet A. *In vivo*, this may be intermolecular resulting in a loop-sheet dimer which extends to form longer polymers (*13*), or intramolecular with a domain-swap of the C-terminal region providing the inter-subunit linkage (*14*). The resulting polymer is deposited within hepatocytes.

The aim of our work was to develop a small molecule corrector of Z α_1_-antitrypsin folding that was able to block the formation of polymers within the endoplasmic reticulum of hepatocytes and that was suitable for oral dosing as a potential treatment for α_1_-antitrypsin deficiency. To achieve this we needed to overcome a number of challenges: (i) the drug target is a highly mobile folding intermediate located in the endoplasmic reticulum; (ii) disparity in the size of the interface between a small molecule and the large protein-protein interaction that it is designed to block; (iii) oral dosing greatly restricts suitable chemical space; (iv) as a non-classical drug target, small molecule binders may well not be well-represented in compound screening libraries; (v) the relatively high concentration of circulating monomeric Z α_1_-antitrypsin (~5μM), even in individuals with severe plasma deficiency, represents a high affinity sink for compound, restricting its access to the target in the hepatocyte and requiring high total blood concentrations of drug to achieve sufficient free drug concentration and target engagement in the liver.

## Results

### Identification of GSK716 through Encoded Library Technology screening, structure guided drug design and cellular profiling

Z α_1_-antitrypsin is a conformationally dynamic molecule (*7, 15*) that polymerises from a near-native conformation late in the folding pathway (*16, 17*) and therefore represents a non-classical target for drug discovery. A cell-free assay approach to hit finding was undertaken so as not to miss compounds that bind α_1_-antitrypsin and block polymerisation but lack the molecular properties to cross cell membranes. This comprised of: (i) an Encoded Library Technology (ELT) screen (*18*) of a library comprising nominal diversity of 2×10^12^ unique components to identify binders to Z α_1_-antitrypsin and (ii) a high throughput screen (HTS) of the GSK compound collection (~1.7 million compounds) for small molecules that could block polymerisation of Z α_1_-antitrypsin. In both screening approaches glycosylated Z α_1_-antitrypsin, purified from the plasma of Z α_1_-antitrypsin homozygotes (*19*), was used since this represents the disease-relevant human pathophysiological drug target that populates an intermediate on the polymerisation pathway (*15, 20*). ELT selections were performed by incubating Z α_1_-antitrypsin with DNA-encoded compound libraries for 1 hour at 4°C and 37°C for 3 rounds of selection with subsequent capture of Z α_1_-antitrypsin using α_1_-antitrypsin Select Resin (GE Healthcare). A variation on this protocol using pre-immobilised Z α_1_-antitrypsin was also used for library selections. In the HTS assay, polymerisation of purified Z α_1_-antitrypsin was induced by incubation at 37°C for 72 hours in the presence of test compounds, with end-point quantification of polymers performed using the polymer-specific monoclonal antibody, 2C1 (*21*) in a TR-FRET-based immunoassay. A number of small molecules that could block polymerisation of Z α_1_-antitrypsin were obtained through the HTS but none progressed beyond the early lead optimisation stage. However, a single lead series of chiral hydroxy-carboxamides (GSK425) was identified from the ELT screen that also demonstrated functional activity at blocking polymerisation in the TR-FRET immunoassay (pIC50 6.5) (Fig. 1a, b).

**Figure 1.**
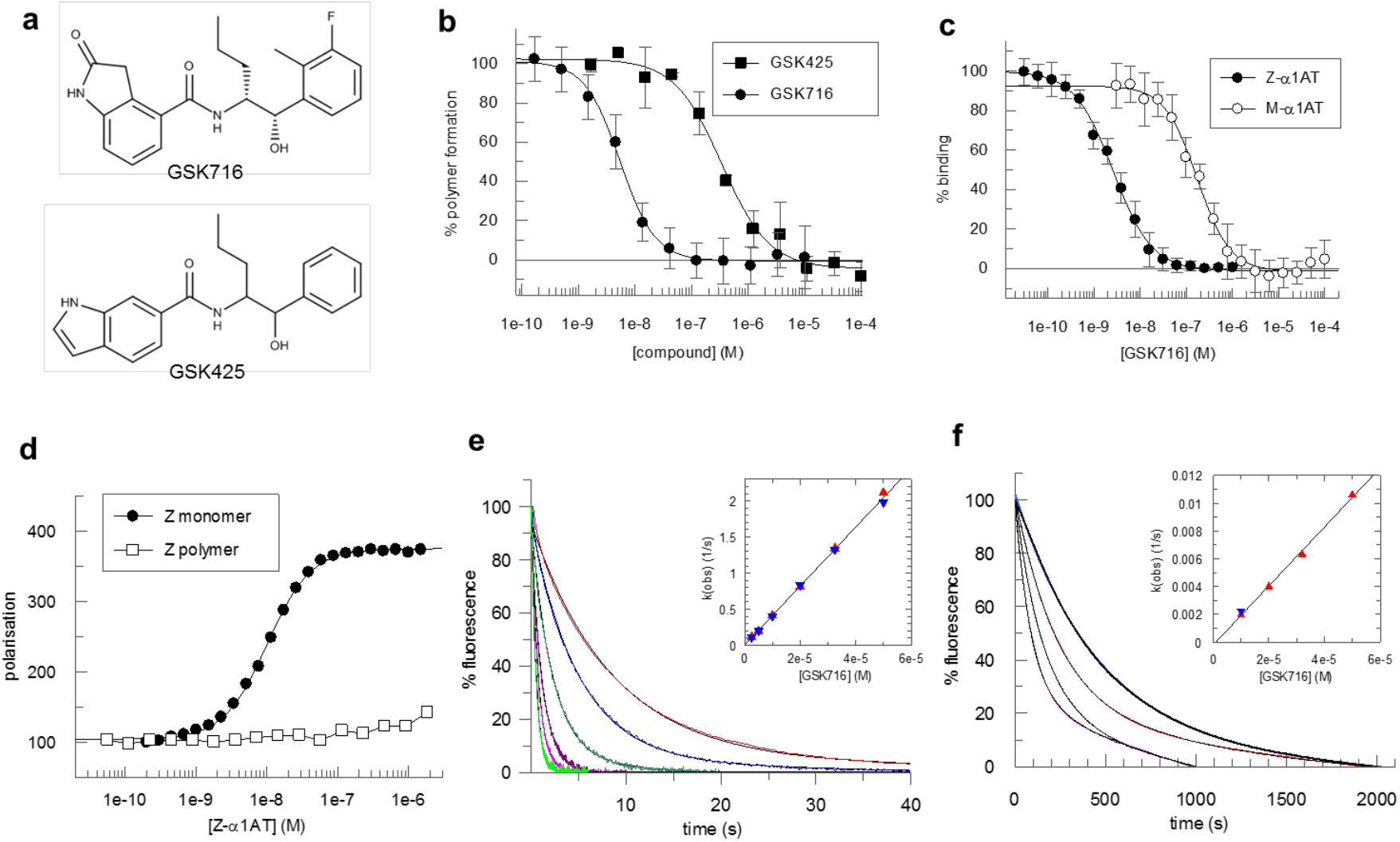
Characteristics of the lead series of chiral hydroxy-carboxamides identified from the ELT screen. (a) The structure of GSK425, identified from the ELT screen, and the derived compound GSK716 obtained through a structure-based design pipeline. (b) The degree of polymerisation of Z α_1_-antitrypsin after 72 hours at 37°C, as determined by an end-point immunoassay using the 2C1 monoclonal antibody, in varying concentrations of compound (Fig 1a and b). Modification of the phenyl and indole heterocycle of GSK425 (pIC50 6.5) resulted in an ~100 fold increase in potency and the discovery of the 2-oxindole GSK716 (pIC50 8.3). (c) GSK716 binds to Z α_1_-antitrypsin with a high affinity mean pKD of 8.5 ± 0.12 (n = 18) as determined by a competition binding assay with a fluorescently labelled derivative. There was a 50-fold lower affinity for plasma-purified wild-type M α_1_-antitrypsin, with a mean pKD of 6.8 ± 0.18 (n = 10). (d) The compound bound to monomeric but not polymeric Z α_1_-antitrypsin (Z α_1_-AT) as reported by fluorescence polarisation of an Alexa-488-labelled variant of GSK716. (e) Representative curves reporting the interaction of different concentrations of GSK716 with Z α_1_-antitrypsin based on changes in intrinsic tryptophan fluorescence. Based on the concentration dependence (*inset*), the second-order rate constant of association was found to be 4.1 x 10^4^ M^−1^ s^−1^. (f) The association of GSK716 with M α_1_-antitrypsin, giving a second-order rate constant of 2.1 x 10^2^ M^−1^ s^−1^.

Optimisation of this initial hit followed a structure-based design approach, exploiting knowledge from iterative crystal structures of small molecule ligands complexed with α_1_-antitrypsin. The central hydroxy carboxamide and propyl chain were found to be critical for binding to Z α_1_-antitrypsin and hence further medicinal chemistry development focussed on modification of the phenyl and indole heterocycle. This resulted in an ~100-fold increase in potency and the discovery of the 2-oxindole GSK716 (pIC50 8.3) (Fig. 1a, b).

### GSK716 is a potent inhibitor of polymerisation *in vitro* and in cell models of disease

GSK716 binds to Z α_1_-antitrypsin with a high affinity mean pKD 8.5 ± 0.12 (n = 18) as determined by a competition binding assay with a fluorescently labelled derivative (Fig. 1c). The binding demonstrates selectivity with a 50-fold lower affinity for plasma-purified wild-type M α_1_-antitrypsin at mean pKD 6.8 ± 0.18 (n = 10) (Fig. 1c). The shape of the curves and native mass spectrometry (not shown) are consistent with a single high-affinity compound binding site. No binding of the fluorescent derivative to polymers of Z α_1_-antitrypsin was observed, indicating conformational selectivity for the monomeric protein (Fig. 1d). The rate of interaction of the compound with the target was monitored through changes in intrinsic tryptophan fluorescence (*10*); this property was used to determine the second-order association rate constants for GSK716 binding to Z (4.1 x 10^4^ M^−1^ s^−1^) and M α_1_-antitrypsin (2.1 x 10^2^ M^−1^ s^−1^) (Figs. 1e, f). From the association rate constants and the affinity values, first-order dissociation rate constants were calculated and found to be of the same order of magnitude for Z (6.1 x 10^−5^ s^−1^) and M α_1_-antitrypsin (1.6 x 10^−5^ s^−1^). Therefore, the selectivity of the compound for Z over M α_1_-antitrypsin is dominated by the difference in the rate of association rather than dissociation.

The ability of GSK716 to block Z α_1_-antitrypsin polymerisation in the ER during folding was assessed by adding GSK716 to CHO-TET-ON-Z-A1AT cells (*8*) with simultaneous induction of Z α_1_-antitrypsin expression using doxycycline. In comparison with controls, GSK716 completely blocked the intracellular formation of Z α_1_-antitrypsin polymers, as measured by staining with the 2C1 anti-Z α_1_-antitrypsin polymer monoclonal antibody (pIC50 = 6.3) (Figs. 2a, b). It also increased the secretion of Z α_1_-antitrypsin approximately 3-fold compared to vehicle control (mean pEC50 6.2 ± 0.23; n = 74) (Fig. 2b). Similar potency between the effects on secretion and polymerisation was observed throughout members of the lead series supporting the hypothesis that these effects are caused by the same pharmacological mode of action. GSK716 had a similar effect on the secretion and polymerisation of constitutively expressed Z α_1_-antitrypsin in iPSC-derived human hepatocytes with the ZZ α_1_-antitrypsin genotype (*22*). It inhibited polymerisation and increased secretion with a mean pIC50 of 6.4 ± 0.45 (n = 16) and mean pEC50 of 6.5 ± 0.37 (n = 14), respectively, inducing an approximately 3-fold increase of secreted levels of Z α_1_-antitrypsin (Figs 2c, d). GSK716 treatment reduced the levels of intracellular Z α_1_-antitrypsin polymer compared with cells assessed before compound addition (Fig. 2c), demonstrating that polymers can be cleared over the time course of the experiment, and that accumulation of polymers is reversible in ZZ-iPSC-hepatocytes.

**Figure 2.**
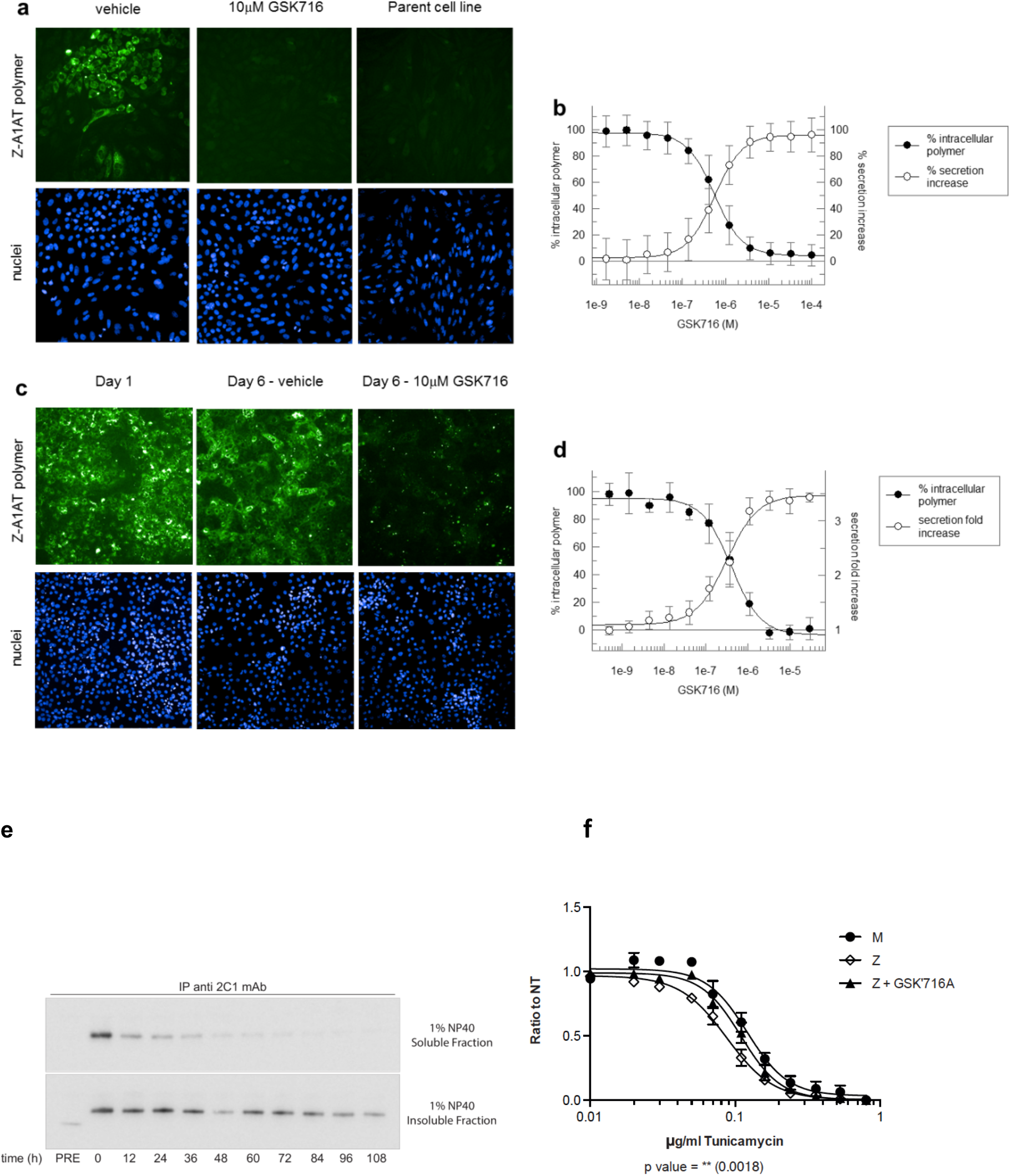
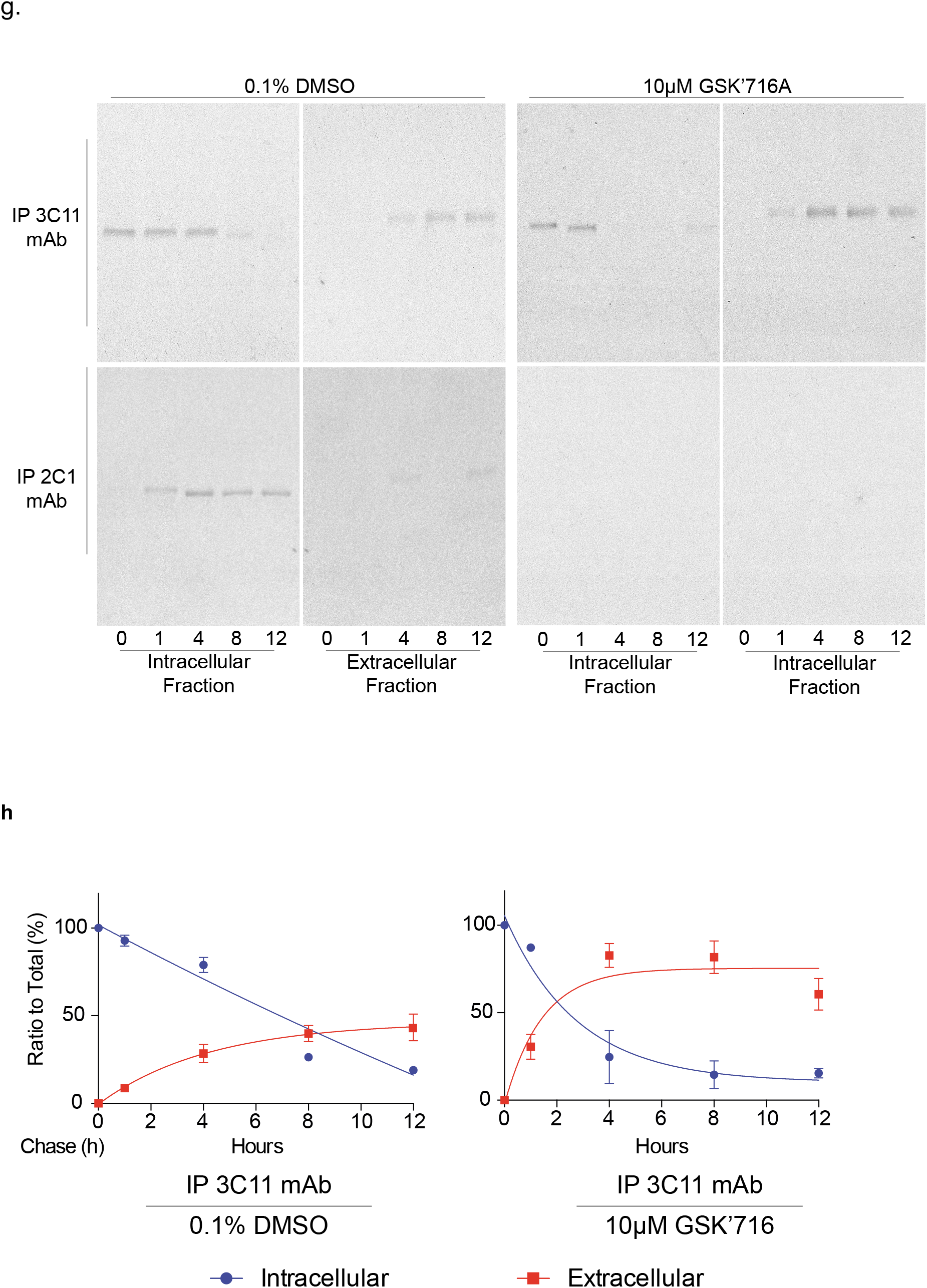
GSK716 inhibits polymerisation of Z α_1_-antitrypsin in cell models of disease. (a) GSK716 was added to CHO-TET-ON-Z-A1AT cells (*8*) with simultaneous induction of Z α_1_-antitrypsin expression using doxycycline, and polymer load was quantified with the 2C1 monoclonal antibody that is specific to pathological polymers of α_1_-antitrypsin (*21*). The parent cell line that did not express Z α_1_-antitrypsin provided a negative control. (b) Quantification of immunostained cells showed that GSK716 completely prevented intracellular polymer formation and increased the secretion of Z α_1_-antitrypsin in a dose-dependent manner. (c) GSK716 was administered to iPSC-derived-hepatocytes and (d) inhibited polymerisation and increased secretion with a similar potency. It induced an approximately 3-fold increase in secreted levels of Z α_1_-antitrypsin. This was apparent even after polymers had been allowed to form. (e) CHO tetracycline-inducible cells expressing Z α_1_-antitrypsin were induced with 0.5 μg/mL doxycycline (Dox) for 48h and treated with 10μM GSK716. Cells are then lysed in 1% v/v NP-40 buffer at different time points (0, 12, 24, 36, 48, 60, 72, 84, 96 and 108 hours). For every time point, NP-40-soluble and insoluble fractions were separated and immunoprecipitated with the 2C1 mAb and separated by 4-12% w/v SDS-PAGE and α_1_-antitrypsin detected by immunoblotting. PRE indicates cells pretreated for 48 hours with 10μM GSK716 and induced for the same time with 0.5 μg/mL doxycycline. The rate of clearance of soluble and insoluble polymer is shown. (f) CHO inducible cells expressing either wildtype M or Z α_1_-antitrypsin (upper panel) were induced with 0.5 μg/mL doxycycline (Dox) and treated with 10μM GSK716 or with 0.1% DMSO (vehicle). After induction for 48 h cells were treated with decreasing doses of tunicamycin as indicated in graph (0.8μg/ml to 0 – 1:1.5 dilution) for 36h. Cell viability was measured by Cell Counting Kit-8. Graph shows mean±SE (M n=5, Z n=8, independent experiments). (g) CHO-K1 Tet-On cells expressing Z α_1_-antitrypsin were induced with doxycycline (0.5μg/ml) for 48 h. Cells subjected to treatment with the experimental compound were incubated with 10μM GSK716 (0.1% v/v DMSO for the control) during the induction. Culture media containing either the experimental compound or the DMSO were changed every 24 h. After the induction, cells were labelled for 10 minutes with ^35^S Met/Cys and chased at the indicated times. Culture media were collected and cells lysed in 1% v/v NP-40 buffer. Intracellular fractions and culture media from cells expressing Z α_1_-antitrypsin were immunoprecipitated either with a mAb against total α_1_-antitrypsin (3C11) or with a polymer-specific mAb (2C1). Samples were resolved by 4-12% w/v acrylamide SDS-PAGE and detected by autoradiography. (h) The graphs show the effect of GSK716 on intracellular and extracellular Z α_1_-antitrypsin (mean ± standard error n=2).

The pre-treatment of CHO cells induced to express Z α_1_-antitrypsin with GSK716 significantly reduced the formation of soluble and insoluble polymers (Fig. 2e compare PRE with time 0). To investigate the ability of GSK716 to protect cells from sensitisation to a secondary insult, Z α_1_-antitrypsin expression was induced in CHO-TET-ON-Z-A1AT cells in the presence or absence of 10 μM GSK716 before exposure to increasing concentrations of the ER stressor tunicamycin (*8*). Cells expressing wildtype M α_1_-antitrypsin were less susceptible to tunicamycin toxicity than cells expressing Z α_1_-antitrypsin in a cell viability assay (Fig. 2f). GSK716 restored sensitivity of Z α_1_-antitrypsin expressing cells to that of the wild-type control cells. The effect of GSK716 on Z α_1_-antitrypsin was confirmed in pulse chase experiments (Fig. 2g and 2h). The small molecule completely blocks the intracellular polymerisation of Z α_1_-antitrypsin and increases secretion of the monomeric protein.

### GSK716 binds to a novel cryptic binding site

A high-resolution crystal structure of α_1_-antitrypsin complexed with the lead compound GSK716 was generated by soaking compound into apo α_1_-antitrypsin crystals (Table 1). The structure reveals that interaction with the compound induces the formation of a cryptic binding site that is not evident in apo structures, at the top of β-sheet-A behind strand 5. This region is referred to as the ‘breach’ as it is the point at which the reactive centre loop first inserts during protease inhibition (*23*), and includes the site of the Z (Glu342Lys) mutation (Fig. 3a). The structure reveals that the 2-oxindole ring of GSK716 stacks with the side chain of Trp194 whilst the carbonyl group forms a hydrogen bond with the mainchain Trp194 (Fig. 3b). Trp194 adopted a new position due to rearrangement of residues Gly192 to Thr203 consistent with the change in intrinsic tryptophan fluorescence induced by binding (Fig. 3c). The phenyl ring and the propyl chain occupy two highly hydrophobic pockets (Figs. 3d, e). Hydrogen bonds are formed between the GSK716 hydroxyl group and the Leu291 backbone, the amide nitrogen hydrogen and the backbone carbonyl oxygen of Pro289, and between the amide carbonyl and the Tyr 244 hydroxyl group (Fig. 3b). This causes displacement of residues Thr339 to Ser 359 of strand 5A relative to the apo protein. Few changes are seen outside of these regions.

**Table 1.** Data collection and refinement statistics

| GSK716 |  |
| --- | --- |
| **Data collection** |  |
| Wavelength | 0.9763 |
| Space group | C2 |
| Cell dimensions |  |
| *a*, *b*, *c* (Å) | 113.954, 39.592 90.517 |
| $\alpha$, $\beta$, $\gamma$ (°) | 90.0, 104.9, 90.0 |
| Resolution (Å) |  |
| *R*<sub>merge</sub> | 0.029 (0.500) |
| CC-half | 0.999 (0.805) |
| *I* / σ*I* | 19.6 (2.2) |
| Completeness (%) | 99.2 (99.2) |
| Multiplicity | 3.3 (3.3) |
| Wilson B-factor | 32.95 |
| **Refinement** |  |
| Resolution (Å) | 55.05-1.76 (1.82-1.76)* |
| No. reflections | 128139 (18557) |
| Uniq reflections | 38875 (5623) |
| *R*<sub>work</sub> / *R*<sub>free</sub> | 18.9 / 22.16 |
| No. atoms |  |
| Protein | 2868 |
| Water |  |
| *B*-factors |  |
| Protein |  |
| Water |  |
| R.m.s. deviations |  |
| Bond lengths (Å) | 0.010 |
| Bond angles (°) | 1.20 |

**Figure 3.**
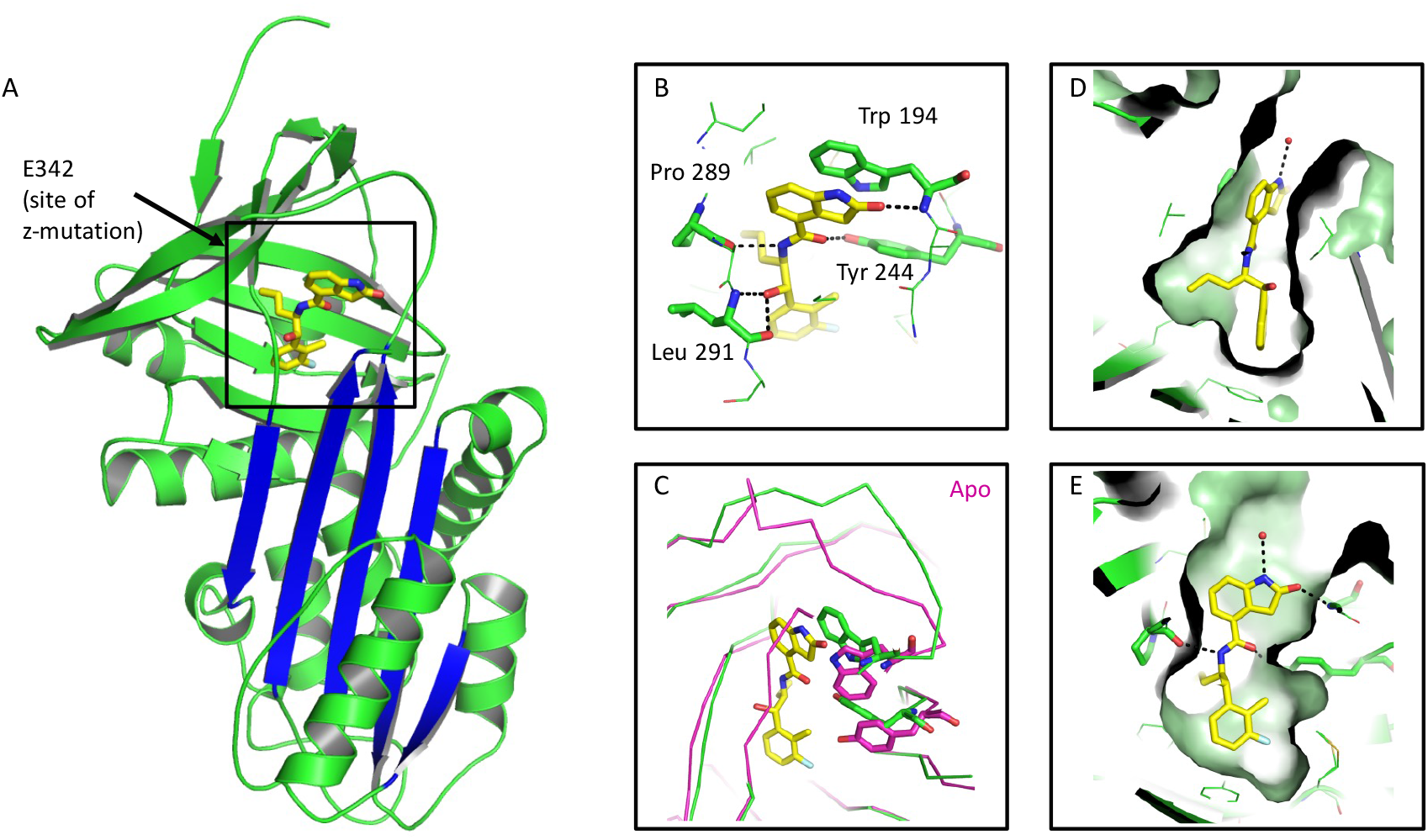
GSK716 binds to a novel cryptic binding site. (a) Cartoon representation of α_1_-antitrypsin with GSK716 shown in yellow stick format. The five-stranded β-A sheet is in blue. (b) Interactions of GSK716 within the cryptic site with key residues shown in stick format and hydrogen bonds in black dashed lines. (c) Overlay of ribbon representation of protein chains of GSK716 complex (green) with apo protein (purple; PDB id 2QUG (*42*)) showing the altered position of Trp194 (stick format) that induces a change in conformation of the Gly192-to-Thr203 loop and re-orientation of Tyr244 (stick format). (d) Surface representation of α_1_-antitrypsin to show propyl and phenyl pocket. (e) Surface representation of α_1_-antitrypsin to show protein-ligand complement of the substituted phenyl and 2-oxindole rings.

### GSK716 interferes with the transition through the polymerisation-prone intermediate M* by stabilising β-sheet A

Polymerisation of α_1_-antitrypsin involves transition through a transient intermediate state known as M*, that is readily populated by the Z variant (*10, 15*). The M* conformational ensemble appears to be a distinct species between the native state conformation and that of the final polymer. One of the hallmarks of M* is its recognition by environment-sensitive fluorescent reporter dyes. Thermal shift assays that make use of the dye SYPRO Orange report the stability of the protein native state against heat-induced unfolding. Experiments, performed using different temperature gradients in the presence and absence of 50μM GSK716 demonstrated a marked increase in the transition midpoint temperature (Fig. 4A); consistent with the stabilization of either or both of the ground- and M*-states of α_1_-antitrypsin (*24*). Correspondingly, in a constant-temperature experiment, oligomers were generated at higher temperatures in the presence of the compound than in its absence when visualised by non-denaturing PAGE (Fig. 4B). Native state stability can also be probed by equilibrium unfolding using chemical denaturants, where a peak in bis-ANS fluorescence corresponds with a maximally populated unfolding intermediate (*25*). The profiles in Fig. 4C show that for guanidinium hydrochloride-induced unfolding this point occurs at a considerably higher denaturant concentration (~1.9M) in the presence of 50 μM GSK716, than in its absence (~1.3M), reflecting an increase in the stability of the native-like state with respect to an unfolding intermediate. The association of GSK716 induced a marked quenching and blue-shift of the α_1_-antitrypsin intrinsic tryptophan fluorescence spectrum (Fig. 4D inset). The rate of change in fluorescence following association of 10μM GSK716 was proportional to the propensity of α_1_-antitrypsin mutants to form polymers *in vivo*: inert (M), mild (S), moderate (Baghdad) and severe (Z) α_1_-antitrypsin.

**Figure 4.**
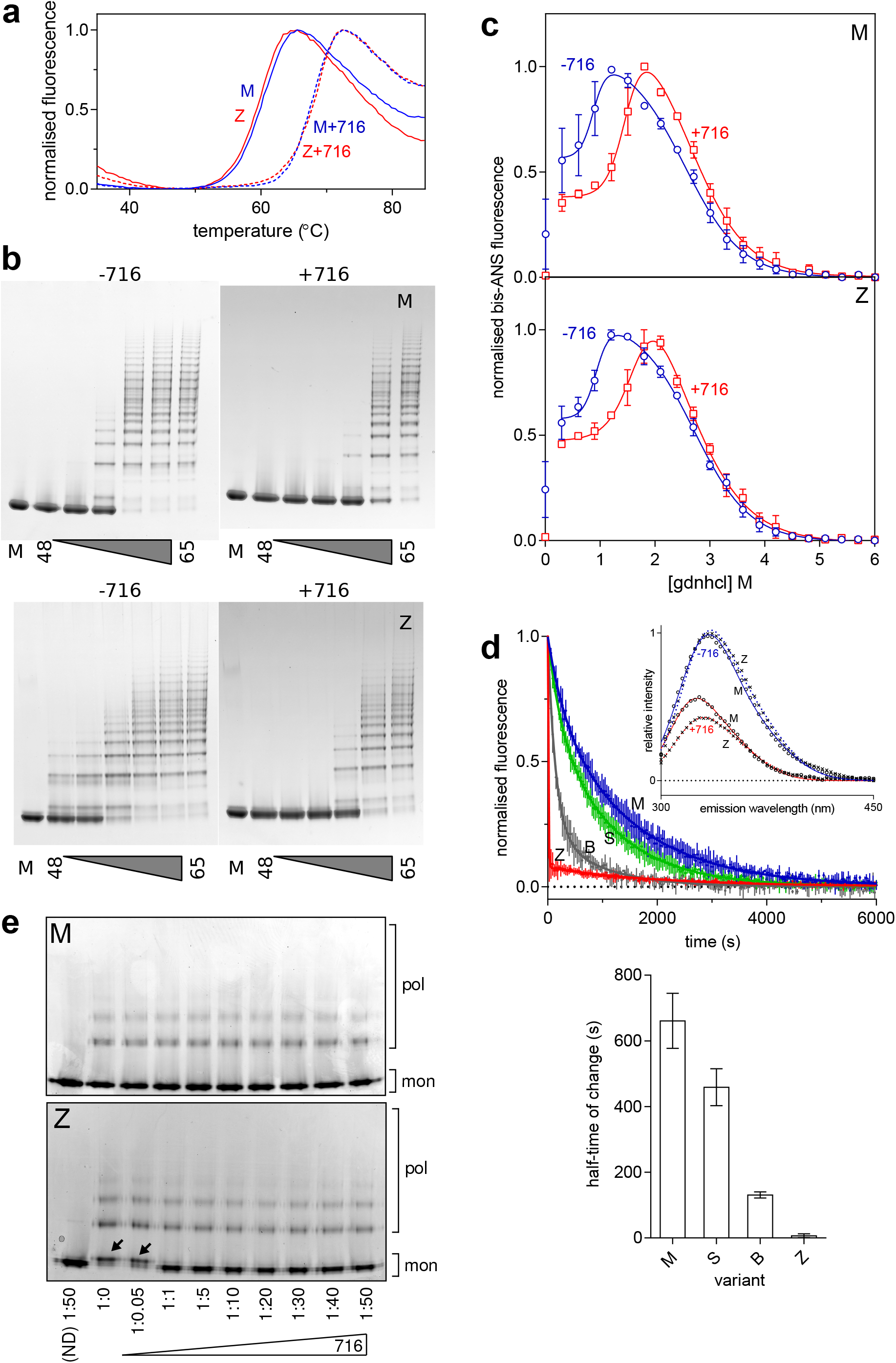
Stabilisation of α_1_-antitrypsin by GSK716. (a) A SYPRO Orange-based thermal stability assay, which reports the transition of M and Z α_1_-antitrypsin from the native to an intermediate state over a 1°C/min thermal ramp, in the presence and absence of 50μM GSK716. (b) M (above) and Z (below) α_1_-antitrypsin, at a concentration of 0.2mg/ml in PBS+5% v/v glycerol, were heated at a range of temperatures between 48°C-65°C for 4 hours in the presence and absence of GSK716, as indicated. The oligomerisation state was determined by non-denaturing PAGE electrophoresis; the lane denoted ‘M’ contains the unheated monomeric control. (c) M and Z α_1_-antitrypsin were subjected to equilibrium unfolding into different concentrations of guanidine hydrochloride (gdnhcl) in the presence and absence of GSK716, with bis-ANS dye added to report the presence of the unfolded intermediate. The normalised fluorescence intensity data were fitted with an equation describing a three-state unfolding curve. Values shown are the mean of three independent experiments and the error bars represent ±SEM. (d) *Top panel*, the association of GSK716 induced a marked quenching and blue-shift of the α_1_-antitrypsin intrinsic tryptophan fluorescence spectrum with respect to unbound (*inset graph*). The association of 10μM GSK716 with four α_1_-antitrypsin variants that vary in their propensity to polymerise, ranging from inert (M), to mild (S), moderate (Baghdad denoted ‘B’) and severe (Z). Representative progress curves of the change in intrinsic tryptophan fluorescence at 330nm for 0.2mg/ml protein are shown. *Bottom panel*, the half-time of association calculated from three such independent experiments (error bars are ±SEM) show a correspondence with the polymerisation propensity of the four variants. The change in Z α_1_-antitrypsin fluorescence was faster than the dead-time of the apparatus (~10s) and scaling was with reference to unbound intensity. (e) Nondenaturing PAGE characterisation of the conformational state of M and Z α_1_-antitrypsin, after rapidly refolding by snap dilution from 6M urea in the presence or absence of GSK716 at the molar ratios indicated. The migration of polymers (pol) and monomers (mon) are shown. Arrows indicate the misfolded by-product M* arising at sub-stoichiometric concentrations of compound.

To investigate whether this activity was consistent with action as a chemical chaperone, M and Z α_1_-antitrypsin were unfolded *in vitro* into 6 M guanidine hydrochloride, and rapidly refolded by snap dilution into denaturant-free buffer in the presence or absence of GSK716. Electrophoresis of the products by non-denaturing PAGE showed an anodally-shifted migration for the Z variant in the absence of compound (Fig. 4E) consistent with a misfolded by-product of M* (*13*), which was corrected at stoichiometric concentrations and above.

Mutations that interfere with the opening of β-sheet A or that perturb its interaction with the N-terminal portion of the reactive centre loop alter the ability of serpins to inhibit target proteases (*24, 26*). The stoichiometry of inhibition (SI) was determined for M and Z α_1_-antitrypsin in discontinuous experiments against a model target protease, chymotrypsin. The pre-incubation of both variants with GSK716 led to a >98% loss of protease inhibitory activity (Fig. 5A). Resolution of the products of the interaction by SDS-PAGE showed full cleavage of the reactive centre loop (Fig. 5B); therefore, this is not a consequence of the protease recognition site in the reactive centre loop becoming inaccessible to the enzyme. These data are consistent with a mechanism in which the compound stabilises β-sheet A against conformational change that mediates both inhibitory activity and pathological misfolding.

**Figure 5.**
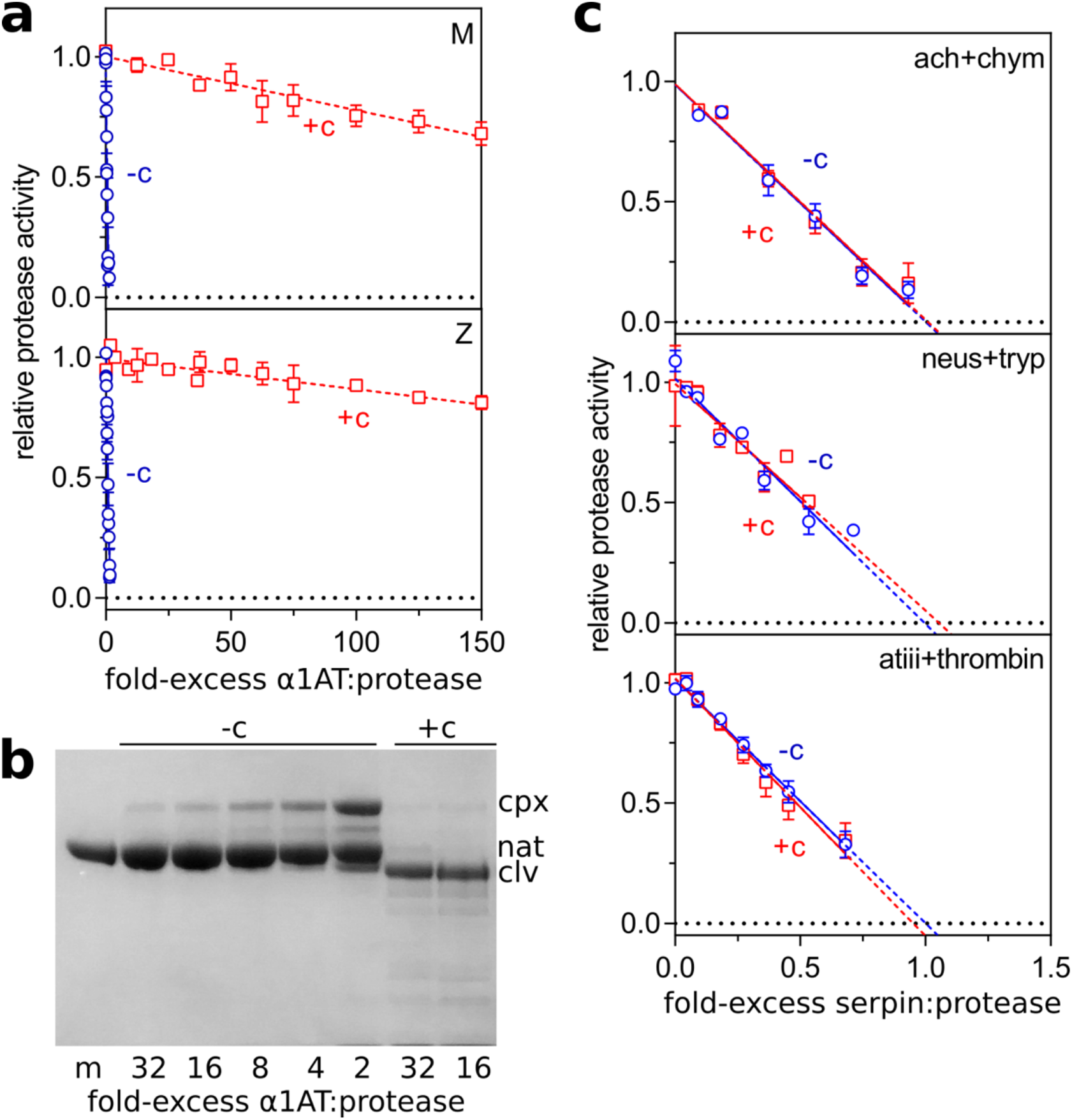
The effect of GSK716 on serpin inhibitory activity. (a) α_1_-Antitrypsin was incubated at varying molar ratios with the model protease bovine α-chymotrypsin, and the residual protease activity determined. The intercept of the regression with the abscissa reflects the number of molecules of α_1_-antitrypsin required to inhibit one molecule of chymotrypsin in the presence (+c) and absence (-c) of 50μM GSK716. (b) M α_1_-antitrypsin was incubated with different molar ratios of chymotrypsin and resolved by SDS-PAGE. The position of covalent α_1_-antitrypsin-chymotrypsin complex (cpx), native (nat) and cleaved (clv) is shown. (c) The inhibitory activity of α_1_-antichymotrypsin against chymotrypsin (ach+chym), neuroserpin against trypsin (neus+tryp), and antithrombin against thrombin (atiii+thrombin) was determined in the presence and absence of GSK716. Error bars reflect ±SD of two independent experiments.

### Characterisation of drug-like properties of GSK716

GSK716 selectivity and PK properties were profiled in order to investigate the suitability of GSK716 for progression into *in vivo* studies and the potential for taking it forward as a clinical candidate for testing in humans. Since GSK716 results in loss of inhibitory activity of α_1_-antitrypsin, the effect of the compound was assessed on other closely-related serpins. GSK716 did not effect the inhibitory activity of antithrombin, neuroserpin and α_1_-antichymotrypsin towards their cognate proteases (Fig. 5C). Furthermore, there were no off target effects in a panel of assays considered predictive of known safety liabilities that precluded further development of GSK716 (Suppl. Table 1).

Since GSK716 exhibited a good level of selectivity over the off-target panel and over other serpins, we determined the *in vitro* and *in vivo* PK properties of the molecule with a view to exploring target engagement *in vivo*. GSK716 has a measured ChromLogD (pH7.4) of 3.8, low binding to human serum albumin (84.2%) and good solubility of amorphous drug substance in FaSSIF (969μg/ml). Permeability in MDR1-MDCK cells in the presence of pgp inhibitor GF120918 was high at 248 and 240 nm.s^−1^ for the apical to basal and basal to apical directions respectively. (N53531-23 for DI). GSK716 exhibited low metabolic clearance in human hepatocytes (0.31 ml/min/g tissue), with moderate to high clearance in mouse hepatocytes (4.56 ml/min/g tissue). It exhibited weak time dependent inhibition of CYP3A4 resulting in a 1.59 fold shift in IC50. Taken together, oral bioavailability is predicted to be high in human, with measured F in rat (48%) and dog (71%) at ≤3mg/kg being consistent with 100% absorption and losses via first pass metabolism only. Mean exposure of GSK716 in blood in the male CD-1 mouse increased with dose following single PO administration at 10, 30 or 100mg/kg (mean dose-normalised C_max_ 58±112, 113±27 and 113±27; DNAUCinf 202±101, 294±47 and 403±246, respectively).

### GSK716 increases secretion of Z α_1_-antitrypsin in a transgenic mouse model of Z α_1_-antitrypsin deficiency

GSK716 was evaluated in a transgenic mouse model with an engineered random insertion of the human Z α_1_-antitrypsin gene (*6*). The PK-PD relationship of GSK716 was explored by dosing Z α_1_-antitrypsin transgenic animals with 10, 30 or 100 mg/kg GSK716 three times a day. Blood and liver were harvested on day 6 at 3hr (~C_max_) and 8hr (C_min_) after the dose for the measurement of total and free drug in both tissues. Blood was also harvested for the measurement of monomeric Z α_1_-antitrypsin in plasma. Total concentrations of GSK716 were determined by LC-MS/MS and the free drug in both tissues was determined using equilibrium dialysis to determine free fraction in the samples, subsequently used to derive unbound concentrations. Blood concentrations demonstrated that the C_min_ levels of free drug were at or above 300nM, the cellular secretion assay EC50, for the majority of the dosing period following 100mg/kg dosing, whereas 30mg/kg and 10mg/kg doses resulted in free drug levels in blood significantly below the cellular EC50 concentrations for a large part of the dosing period. Both free and total drug concentrations of GSK716 at the targeted site of action in the liver were equivalent to those in blood (Table 2).

**Table 2.**
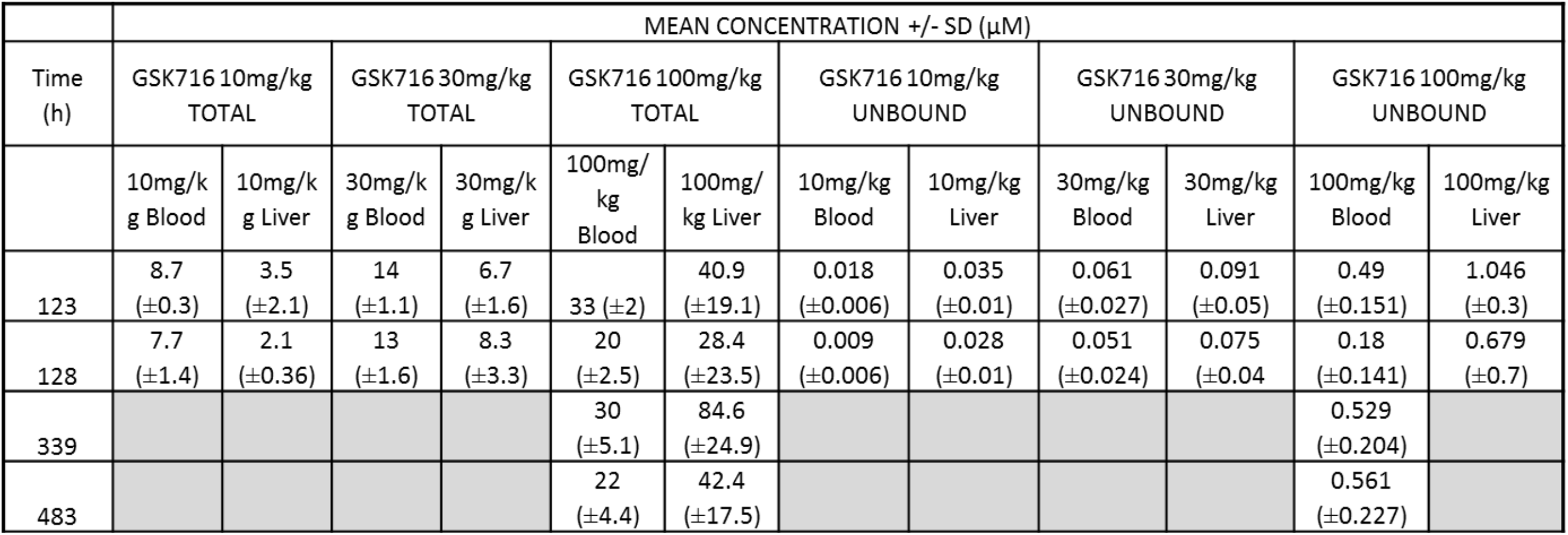
Unbound and total drug concentrations of GSK716 following dosing in Z α_1_-antitrypsin transgenic mice

A significant fraction of the total Z α_1_-antitrypsin in the circulation is in the polymeric conformation (*27*). There are no antibodies that are specific for monomeric Z α_1_-antitrypsin and so to directly determine its concentration, a deconvolution method was developed based on immunoassays with antibodies for either total or polymeric α_1_-antitrypsin, and calibration curves with purified monomeric and polymeric Z α_1_-antitrypsin. Monomeric Z α_1_-antitrypsin was measured in plasma samples following 6 days of dosing and levels were normalised to each animals’ predose control levels to account for the natural variation of Z α_1_-antitrypsin between animals. Administration of 100mg/kg GSK716 resulted in a mean 7-fold increase in circulating monomeric Z α_1_-antitrypsin levels demonstrating robust target engagement in the liver (Fig. 6A). Interestingly, 30mg/kg and 10mg/kg groups also gave significant, dose-dependent increases in circulating Z α_1_-antitrypsin despite free concentrations being below the cellular EC50 for secretion for much or all of the dosing period. Total drug levels and changes in Z α_1_-antitrypsin following 3 days of dosing were indistinguishable to those following 6 days of dosing. There was no effect on circulating serum albumin after 5 days of dosing which provides evidence that GSK716 is specific for Z α_1_-antitrypsin. Moreover the effects are not mediated by metabolites of GSK716 as the major metabolites have much reduced or no binding to α_1_-antitrypsin.

**Figure 6.**
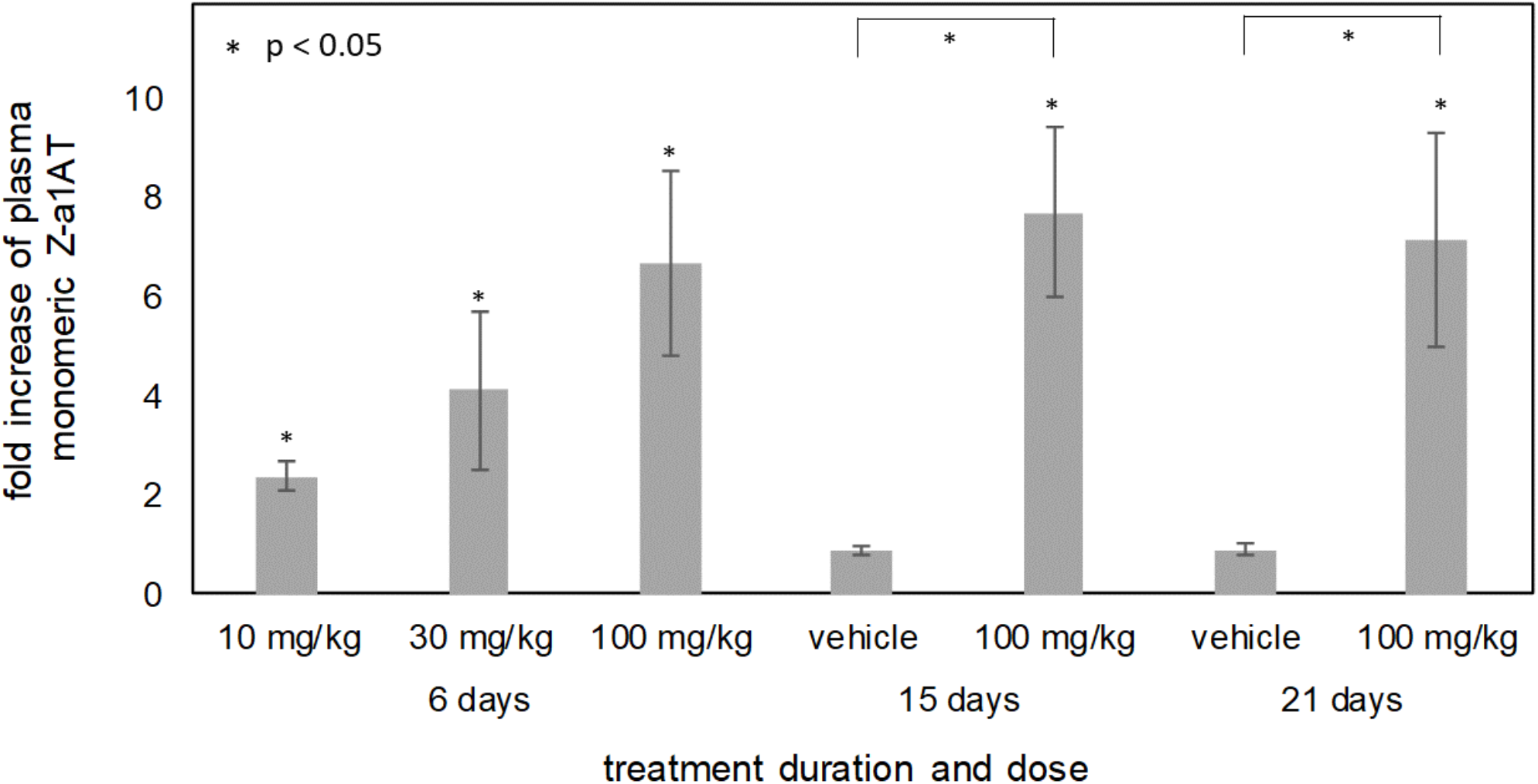
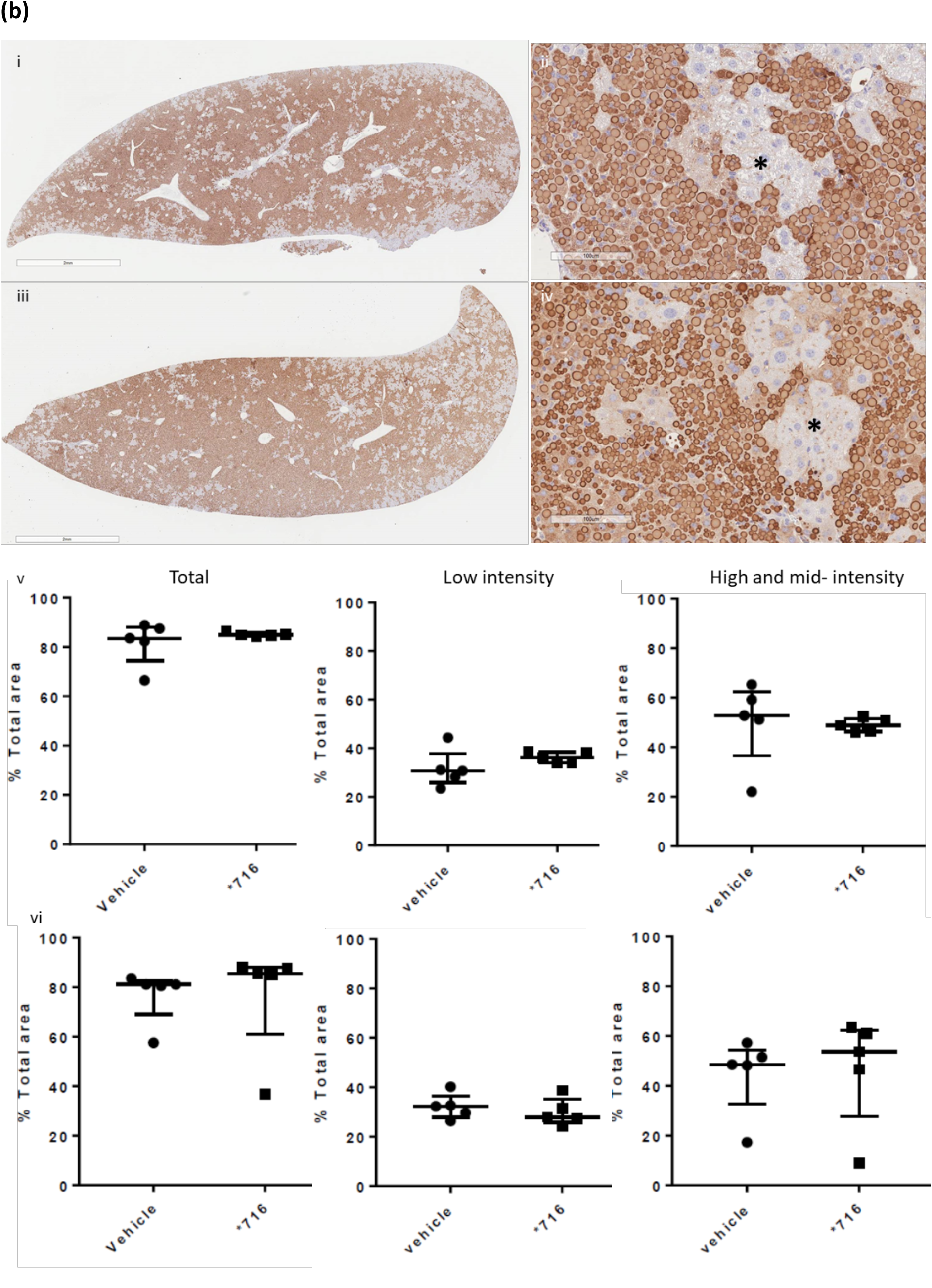
GSK716 increases secretion of Z α_1_-antitrypsin in a transgenic mouse model of Z α_1_-antitrypsin deficiency. Z α_1_-antitrypsin transgenic animals were dosed with 10, 30 or 100 mg/kg GSK716 three times a day. Total concentrations of GSK716 were determined by LC-MS/MS (Table 2). (a) 100mg/kg GSK716 resulted in a mean 7-fold increase in circulating monomeric Z α_1_-antitrypsin levels demonstrating robust target engagement in the liver. The observed steady state total and free compound levels of GSK716 in the transgenic Z α_1_-antitrypsin mouse were predicted well by the *in silico* PK model built on: (i) *in vitro* metabolic clearance data, (ii) plasma protein binding data, (iii) *in vivo* PK data from wild type mice and (iv) a term comprising a 5μM circulating sink for drug with an affinity of 1.5 nM, representing the Z α_1_-antitrypsin within blood. The target free drug concentration was selected based on the observed potency in the *in vitro* secretion assays in which the total drug approximates to the free drug in the assay. Interestingly, 30mg/kg and 10mg/kg groups also gave significant, dose-dependent increases in circulating Z α_1_-antitrypsin despite free concentrations being below the cellular EC50 for secretion for much or all of the dosing period. Significance at p<0.05 by a student’s t test is denoted by *: pairwise: each animal is compared with itself, treated vs pre-treatment; between groups: compound treated vs vehicle. (b) Representative example images of 2C1 stained livers from 20 day vehicle (i and ii) and 100mg/kg TID GSK716 (iii and iv) treated animals. The asterisks indicate regions of hepatocytes that are negative on 2C1 immunostaining. Quantification of 2C1 stained area in livers from vehicle and 100mg/kg TID GSK716 treated animals at day 15 (v) and 21 (vi), shown as total area and low-only or high and mid-only intensity stained areas. There were no significant differences between groups by KW ANOVA with Tukey’s post-hoc analysis, data are the median +/− interquartile ranges.

Since GSK716 blocks polymer formation in cells we explored the effect of dosing GSK716 on liver polymer levels. Z α_1_-antitrypsin polymers formed in CHO-TET-ON-Z-A1AT and ZZ-iPSC-hepatocytes are cleared from cells with a half-life of between 8 and 48 hours depending on whether they partition to the soluble or insoluble fractions (Ronzoni *et al*, manuscript submitted for publication). Since the compound does not bind Z α_1_-antitrypsin polymer, an effect on total liver polymer levels will be dependent on the rate at which the liver can clear the polymer already present and the rate at which polymer continues to accumulate in animals not treated with drug. GSK716 was dosed at 100 mg/kg three times a day for 20 days. Monomeric Z α_1_-antitrypsin increased by a mean of 7-8 fold in plasma samples over the pre-dosing baseline levels on days 15 and 21 of dosing, similar to the effect in animals dosed for 3 or 6 days and consistent with sustained target engagement through the dosing period (Fig. 6a). Liver polymer levels were investigated by staining with 2C1 anti-polymer monoclonal antibody and were scored blinded by a pathologist or by quantification using an algorithm to measure all areas of positive staining (Suppl. Fig. 1). There was no difference observed in total liver polymer load when assessed by manual or quantitative scoring (Fig. 6b). There was no significant fibrosis in any of the liver sections. Intrahepatic α_1_-antitrypsin was also assessed by ELISA. The vast majority (95-100%) of α_1_-antitrypsin within the liver is polymer with the monomer typically being below the level of detection. Treatment with GSK716 increased the monomer measured in liver homogenate by approx. 4-fold in keeping with the changes seen in blood.

## Discussion

Disrupting protein-protein interactions with small molecule compounds that maintain drug-like properties is a significant challenge. Here we describe the identification of a potent and selective α_1_-antitrypsin corrector, GSK716 that abolishes intracellular polymerisation of Z α_1_-antitrypsin and increases the circulating levels of monomeric protein by 7-fold in a transgenic mouse model of disease. The co-crystal structures with α_1_-antitrypsin demonstrate that the small molecule ameliorates the effect of the Glu342Lys (Z) mutation by: (i) optimisation of hydrophobic packing in the breach region; (ii) formation of hydrogen bonds with buried polar atoms; and (iii) displacement of the backbone at the top of strand 5A into a configuration less compatible with partial loop insertion. This movement of strand 5A is an early step in reactive loop-β-sheet A models of polymerisation (*11*) and an obligate one in the C-terminal polymer linkage (*14*). Following GSK716 binding there is a marked stabilisation of α_1_-antitrypsin against the conformational changes associated with M* intermediate formation. This in turn increases folding efficiency thereby reducing the formation of polymers. Precedent for this general mechanism comes from our development of a tool monoclonal antibody that exerted a similar effect on Z α_1_-antitrypsin (*16, 17*).

The GSK716 induced displacement at the top of strand 5A is consistent with the association rate-driven preference for Z α_1_-antitrypsin and an increased availability of the cryptic pocket. The pocket, once formed, appears to be structurally equivalent in both wildtype M and mutant Z α_1_-antitrypsin as reflected by a similar rates of dissociation of GSK716. Binding of GSK716 to α_1_-antitrypsin increased with the propensity of mutants to form polymers: M, S, Baghdad and Z α_1_-antitrypsin, indicating that pocket formation and polymerisation are intimately linked. This mode of action is compatible with the lack of binding of GSK716 to polymers, in which partial or complete insertion of the reactive centre loop completes a β-hairpin turn and so occludes the compound binding site. The finding that GSK716 mediates its action by binding to a cryptic pocket implies that intrahepatic polymers form from a near-native or native conformation, rather than a more extended intermediate.

GSK716 blocks Z α_1_-antitrypsin polymerisation in cell free media and in the ER of both CHO and iPSC models of α_1_-antitrypsin deficiency. It increased secretion from these cell models by approximately 3-fold with similar potency in each model. Treatment with GSK716 reduced the levels of intracellular Z α_1_-antitrypsin polymer compared with cells assessed before compound addition demonstrating that polymers can be cleared over the time course of the experiment, and that accumulation of polymers is reversible in ZZ-iPSC-hepatocytes. These findings were confirmed by pulse chase experiments which showed that GSK716 abolished intracellular polymers (when assessed by the 2C1 mAb) and increased the clearance and secretion of Z α_1_-antitrypsin. Dosing of GSK716 in transgenic mice that express Z α_1_-antitrypsin increased circulating levels of Z α_1_-antitrypsin by 7-fold within 3 days of dosing indicating robust target engagement. This effect was maintained for 21 days. The increase in Z α_1_-antitrypsin in the circulation at 10 and 30mg/kg of GSK716 was surprising given that systemic free drug levels were below the cellular EC50 for secretion for much or all of the dosing period. The reason for this is unclear but it is possible that the target engagement *in vivo* is greater than predicted from the potency in the *in vitro* cellular assays. Alternatively, it is possible that the first pass effect of drug reaching the liver immediately after absorption delivers some efficacy over that predicted from modelling the compound concentration at steady state levels. Together these data suggest potential upsides for the required compound exposure to deliver efficacy in individuals with Z α_1_-antitrypsin deficiency.

Despite the increase in plasma levels of Z α_1_-antitrypsin, there was no reduction in intrahepatic Z α_1_-antitrypsin inclusions after 20 days of dosing in the transgenic mouse. This may be because the Z α_1_-antitrypsin is released from the globule devoid hepatocytes or because the intrahepatic polymers in the transgenic mice are not cleared as readily as those that are generated over a few days in model systems such as CHO cells and iPSC-hepatocytes. Our findings are in contrast to work with the autophagy activator Carbamazapine which had profound effects on liver polymer in Z α_1_-antitrypsin transgenic mice following 2 weeks of dosing (*28*). RNAi approaches that inhibit Z α_1_-antitrypsin expression and polymer formation have reported decreases in Z α_1_-antitrypsin in transgenic mouse liver following 12-33 weeks of treatment, albeit without reports of data at earlier timepoints (*29*). It is likely that GSK716 will need to be dosed to transgenic Z α_1_-antitrypsin mice for significantly longer than 20 days to demonstrate an effect on total liver polymer levels. It remains to be seen whether the intra-hepatic polymer needs to be cleared from the liver in order to have some functional benefit or whether the accumulated polymer inclusions are inert and abrogation of polymer production is sufficient to restore the functioning of the ER and hence the health of the cells. Moreover treatment with GSK716 may be sufficient to protect against the two-hit process whereby the Z α_1_-antitrypsin polymers sensitise the liver to a secondary insult such as alcohol, drug or liver fat (*8*) (*9*).

Polymer formation and inhibitory activity are inextricably linked in the serpin mechanism (*30*) and thus small molecules that block polymerisation may have the unwanted effect of also blocking inhibitory activity. Bound GSK716 inhibits the serpin activity of α_1_-antitrypsin and so would not be expected to increase protease inhibitory activity during the dosing period. However, the half-life of monomeric Z-α_1_-antitrypsin in human is 6 days whereas drug would be expected to be cleared with a half life of a few hours after dosing raising the possibility of a pulsatile dosing regimen that would lead to increased, active serpin. The slow development of the lung disease in individuals with α_1_-antitrypsin deficiency over many decades suggests that acute effects associated with inhibition of serpin activity are unlikely.

There is increasing recognition that heterozygosity for wildtype M and mutant Z α_1_-antitrypsin alleles predisposes to liver disease (*9*). Our small molecule approach to block polymer formation has an advantage over siRNA therapies to ‘knock down’ α_1_-antitrypsin production in treating the MZ α_1_-antitrypsin heterozygote. GSK716 has a 100-fold greater affinity for Z than M α_1_-antitrypsin and so may be dosed at a level that prevents polymerisation of Z α_1_-antitrypsin without reducing the inhibitory effect of the wildtype M protein.

In summary we report the first small molecule drug-like correctors of Z α_1_-antitrypsin folding obtained via optimisation of hits from an Encoded Library Technology screen (*18, 31, 32*) that are suitable for oral delivery, correct folding in human patient iPSC-derived hepatocytes and increase circulating Z α_1_-antitrypsin levels in a transgenic mouse model of α_1_-antitrypsin deficiency.

## Materials and Methods

Alphα_1_-antitrypsin was purified from the plasma of M (wildtype) and Z α_1_-antitrypsin homozygotes and recombinant Cys232Ser α_1_-antitrypsin was expressed and purified as detailed previously (*19, 33, 34*). The data generated in this manuscript have used a number of different preparations of GSK716 all of which have been checked for identity and purity by NMR / MS. Moreover all batches have been tested for activity in blocking α_1_-antitrypsin polymerisation in the assay as shown in Fig 1b. Full experimental procedures and analytical data for synthesising GSK716 are available in patent WO2019/243841A1 (‘compound 1’).

### DNA-encoded library technology (ELT) screen

An encoded library technology (ELT) screen with a nominal diversity of 2×10^12^ unique components was used to identify small molecules that bind monomeric Z α_1_-antitrypsin at 4°C, 22°C, and 37°C. Affinity selections were performed as described previously (*31*).

### *In vitro* assay of Z α_1_-antitrypsin polymerisation

An antibody-based time-resolved fluorescence resonance energy transfer (TR-FRET) assay was developed to monitor the polymerisation of 5nM Z α_1_-antitrypsin following incubation with varying concentrations of compounds at 37°C for 72 hours. This assay used the 2C1 monoclonal antibody (1.25nM) that is specific to pathological polymers of α_1_-antitrypsin (*21*), a polyclonal antibody (1/320 dilution) that binds to all forms of α_1_-antitrypsin (Abcam product 9373), an anti-mouse IgG (1.5nM) labelled with fluorescence donor Eu-W1024 (Perkin Elmer product AD0076) and an anti-rabbit IgG (14.3nM labelled with acceptor (APC). In the presence of polymeric Z α_1_-antitrypsin, a 4-antibody sandwich is formed allowing energy transfer to occur between the Europium- and Allophycocyanin fluorophores. The TR-FRET signal was read on an Envision plate reader (PerkinElmer), by excitation of Europium at 337nm and detection of emission at 665nm and 620nm.

### Compound association experiments

Kinetic parameters of GSK716 binding to M, Z, S (Glu264Val) and Baghdad (Ala336Pro) (*35*) α_1_-antitrypsin were measured by detecting intrinsic tryptophan fluorescence of the protein (excitation at 280 nm and detection of emission at 320 nm) on a stopped flow apparatus (Applied Photophysics) (*10, 36*). A competition assay for binding to M and Z α_1_-antitrypsin and Z α_1_-antitrypsin polymers (*19, 33, 34*), based on an Alexa488-labelled analogue of GSK716 (A488-GSK716), was used to determine the binding affinity of test compounds.

### Thermal stability, unfolding and assessment of protease inhibition

The native state stability of α_1_-antitrypsin on addition of compounds was investigated by thermal denaturation in the presence of a 5X concentration of SYPRO Orange dye solution (Life Technologies) (*37*). Resistance to heat-induced polymerisation was determined using an end-point constant-temperature assay. Equilibrium unfolding was evaluated with a Bis-ANS dye (*10*) and rapid refolding following denaturation in 6M urea was assessed by non-denaturing PAGE. The inhibitory activity of α_1_-antitrypsin was measured by titration against the model protease bovine α-chymotrypsin. The activity of antithrombin, neuroserpin and α_1_-antichymotrypsin were assessed against human thrombin, bovine trypsin and bovine α-chymotrypsin respectively.

### Crystallisation and structure determination

Crystallisation of recombinant Cys232Ser α_1_-antitrypsin was carried out in 2-well MRC crystallisation plates with a Mosquito robot (TTP Labtech) using 100nl protein solution and 100nl well solution. Crystals grew from 22% w/v PEG1500, 0.2M MES pH6.0 and were soaked for 24hrs with 25mM compound (5% v/v DMSO). X-ray diffraction data were collected at Diamond on beamline IO3. Structure solution was carried out by molecular replacement using PHASER (*38*). The model used to solve this structure was a related complex (data not shown) which had been solved by molecular replacement using PDB entry 2QUG as a starting model. Building was carried out using Coot (*39*) and refinement with REFMAC (*40*).

### *In vitro* pharmacokinetics

ChromLogD was measured as previously described (*41*). Permeability in MDR1-MDCK cells with pgp inhibitor, CYP3A4 TDI fold IC50 shift and hepatocyte clearance were performed as detailed in the supplementary material.

### Measurement of monomeric Z α_1_-antitrypsin in plasma and cell biology

Plasma standards of monomeric and polymeric Z α_1_-antitrypsin were prepared and assayed with an antibody mix comprising a 1:160 dilution of rabbit polyclonal anti-α_1_-antitrypsin (Abcam product 9373), 0.23μg/ml Terbium anti-mouse, 4.5μg/ml Alexa488 goat anti-rabbit, 0.17μg/ml mouse anti-total-α_1_-antitrypsin monoclonal 3C11 and 2μg/ml mouse 2C1 anti-polymeric-α_1_-antitrypsin). After a 16 hour incubation at 20°C, TR-FRET detection was performed on an Envision plate reader (Perkin Elmer).

Plots of the FRET ratio (acceptor/donor signal) versus sample dilution result in bell-shaped curves due to the hook effect. The polymer Z α_1_-antitrypsin concentration was determined by comparing the peak positions of these bell-shaped curves for plasma samples with those for polymer calibration samples. The polymer concentration in the plasma sample and the polymer calibration curve were then used to derive the contribution of polymer to the FRET signal in the total α_1_-antitrypsin assay at the signal peak, enabling determination of the concentration of monomeric Z α_1_-antitrypsin.

#### Z α_1_-antitrypsin secretion /polymerisation in CHO-TET-ON-Z-α_1_AT cells

The accumulation and clearance of Z α_1_-antitrypsin-polymer was measured with the 2C1 monoclonal antibody that is specific for polymerised antitrypsin in CHO-TET-ON-Z-α_1_AT cells (*8*) and iPSC-derived hepatocytes generated from a patient with a PiZZ genotype (*22*). Secretion of Z α_1_-antitrypsin was determined by TR-FRET. Cell viability was analysed with cell counting kit-8 (Sigma) after treating the cells with 10μM GSK716 or 0.1% v/v DMSO (vehicle) and an increasing concentration of tunicamycin (0.8 μg/ml to 0.01 μg/ml).

#### Pulse chase experiments

CHO K1 cells were labelled after 48 h induction with 0.5 μg/ml doxocycline. Cells were pulsed (0.45MBq/10^6^ cells) for 10 min with ^35^S Cys/Met (EasyTagTM Express Protein Labelling, Perkin Elmer, Beaconsfield, UK) in DMEM without Cys/Met, and then chased in normal culture medium for 0, 1, 4, 8 and 12 h. Radiolabelled α_1_-antitrypsin was isolated by immunoprecipitation and resolved by SDS-PAGE followed by autoradiography. Densitometric analysis of α_1_-antitrypsin bands was performed with ImageStudioLite (LI-COR Biosciences, USA). Statistical analysis was performed using the GraphPad Prism program (GraphPad Software, La Jolla, CA, USA).

### *In vivo* experiments

#### Pharmacokinetics

Male CD-1 mice, Han Wistar rats or beagle dogs were administered GSK716 as a suspension in 1% w/v aqueous methylcellulose via oral gavage at doses of 1mg/kg (dog) or 10, 30 and 100mg/kg (mouse, rat). Blood samples were taken into EDTA, diluted and mixed with acetonitrile containing internal standard (alprazolam) and centrifuged to precipitate proteins. Aliquots of the resultant supernatant was analysed by LC-MS/MS and concentrations of GSK716 in blood were determined. Noncompartmental pharmacokinetic analysis (NCA) was carried out using Phoenix WinNonLin 6.3 (Certara L.P.). Free drug was measured in *Z* α_1_-antitrypsin transgenic mouse by dialysing tissue samples in rapid equilibrium dialysis cassettes.

#### Z α_1_-antitrypsin transgenic mouse experiments

Hemizygous female Z α_1_-antitrypsin transgenic mice were 13-15 weeks of age at start of study. Drugs were administered via oral gavage three times a day at 8 hour intervals at 10mL/Kg as a suspension in aqueous methylcellulose containing 0.5% w/v Tween 80. In life blood sampling was performed vial tail nick.

#### Histopathology, immunohistochemistry and image analysis

Formalin-fixed, paraffin embedded livers from all mice were sectioned and stained with haematoxylin and eosin (H&E) and periodic acid Schiff (PAS) with diastase to examine tissue quality and the presence of diastase-resistant polymer inclusions respectively. Immunohistochemistry for polymerised α_1_-antitrypsin was performed using 2C1 monoclonal primary antibody and a Mouse-on-Mouse Polymer kit (Abcam AB127055) for detection. Representative sections of liver from each mouse were blindly evaluated by a Board-Certified Veterinary Pathologist and ranked in order of polymer content. A semi-automated image analysis solution was developed to quantify total polymer content (Supp. Fig 1). Thresholds were optimised to separate staining of the polymer inclusions (high and medium intensity) and more diffuse cytoplasmic staining (low intensity).

## Acknowledgements

We are very grateful to Jeff Teckman, Department of Pediatrics, Saint Louis University, MO, USA for providing the transgenic mouse model of α_1_-antitrypsin deficiency. Riccardo Ronzoni and Imran Haq were supported by GlaxoSmithKline. Alistair Jagger is the recipient of a BBSRC CASE studentship. This work was supported by GlaxoSmithKline, the Medical Research Council (UK) (MR/N024842/1) and the UCLH NIHR Biomedical Research Centre. DAL is an NIHR Senior Investigator. Data collection was performed on beamline I03 at the Diamond Light Source and the authors would like to thank the staff for facility access and technical support.

## Author contributions

DAL, AB and ACP designed the programme of work. DAL, ACP, JAI, IU and DSH served on the joint UCL-GSK DPAc project oversight board. JAI, IH, AJ, AD, DF, JPH, MR, JR and MB undertook the biochemical assessment of the compounds, AO, SJM, AD, MR, TJ and RR assessed the cell biology, CC, KJS and MN undertook structural biology studies, CA, SB, JM, AO, ZZ, SS and KL undertook encoded library technology screening, HD, PE, EJ, RT, LT and SW undertook PK, *in vivo* profiling and pathology, AB, ADe, ND, DH and JL undertook medicinal chemistry. All authors reviewed, revised and approved the final manuscript.

## The Paper Explained

### Problem

Intracellular protein aggregation can result in ‘gain of function’ cell toxicity. It has proved challenging to develop small molecules that can stabilise intracellular mutant proteins, prevent self-aggregation and so ameliorate disease. Severe α_1_-antitrypsin deficiency results largely from the Z allele (Glu342Lys) that causes the accumulation of homopolymers of mutant α_1_-antitrypsin within the endoplasmic reticulum of hepatocytes in association with liver disease.

### Results

We have undertaken a medicinal chemistry campaign to develop an orally bioavailable small molecule that binds to intra-endoplasmic reticulum mutant Z α_1_-antitrypsin, corrects the folding defect and increases secretion in a transgenic model of disease.

### Impact

This study reports the successful targeting of an aggregation-prone mutant in order to prevent the intracellular polymerisation and accumulation of α_1_-antitrypsin that underlies α_1_-antitrypsin deficiency. It provides proof-of-principle that the identification of ‘mutation ameliorating’ small molecules is a viable approach to treat protein conformational diseases.

## Supplementary results

**Supplementary Table 1.**
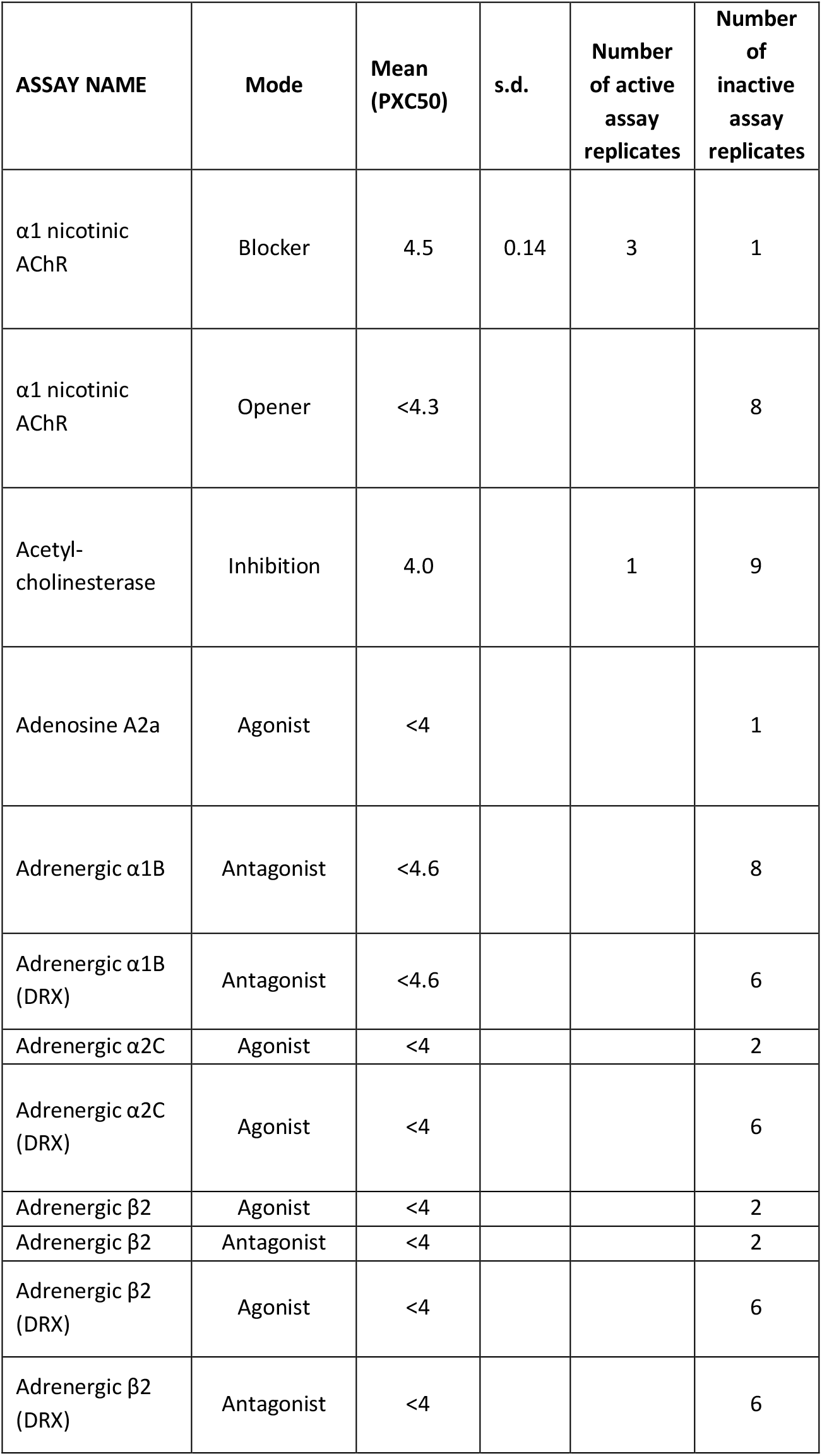
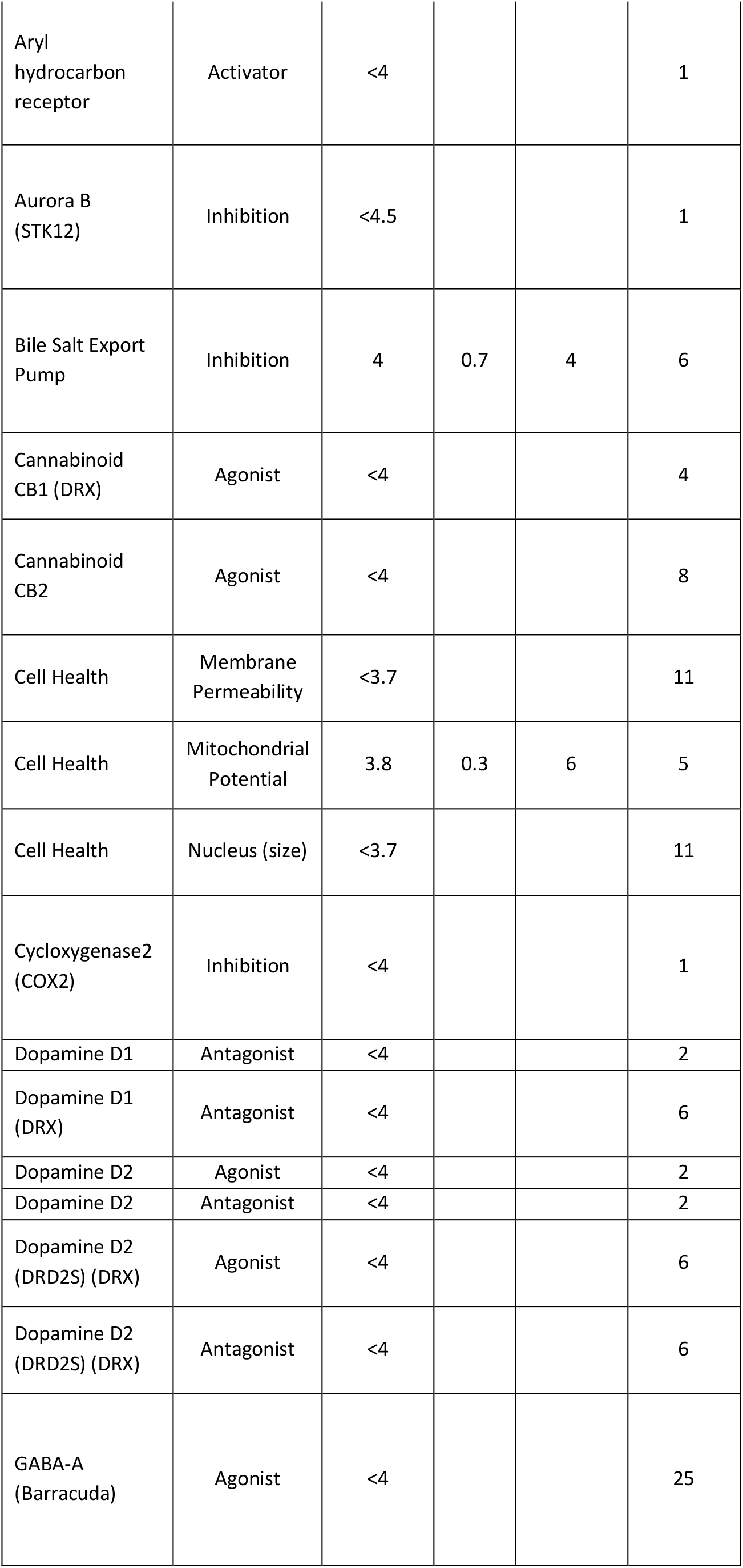
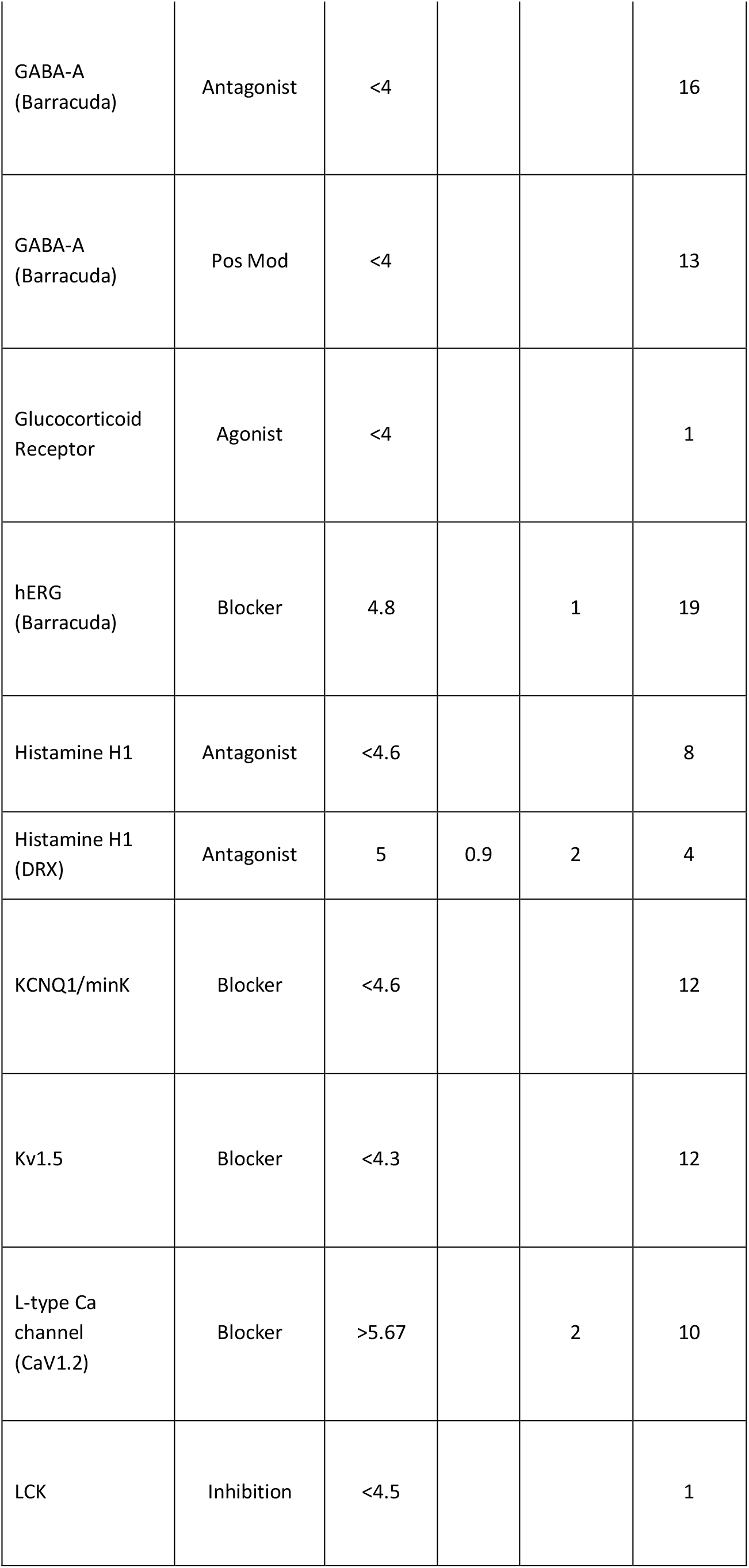
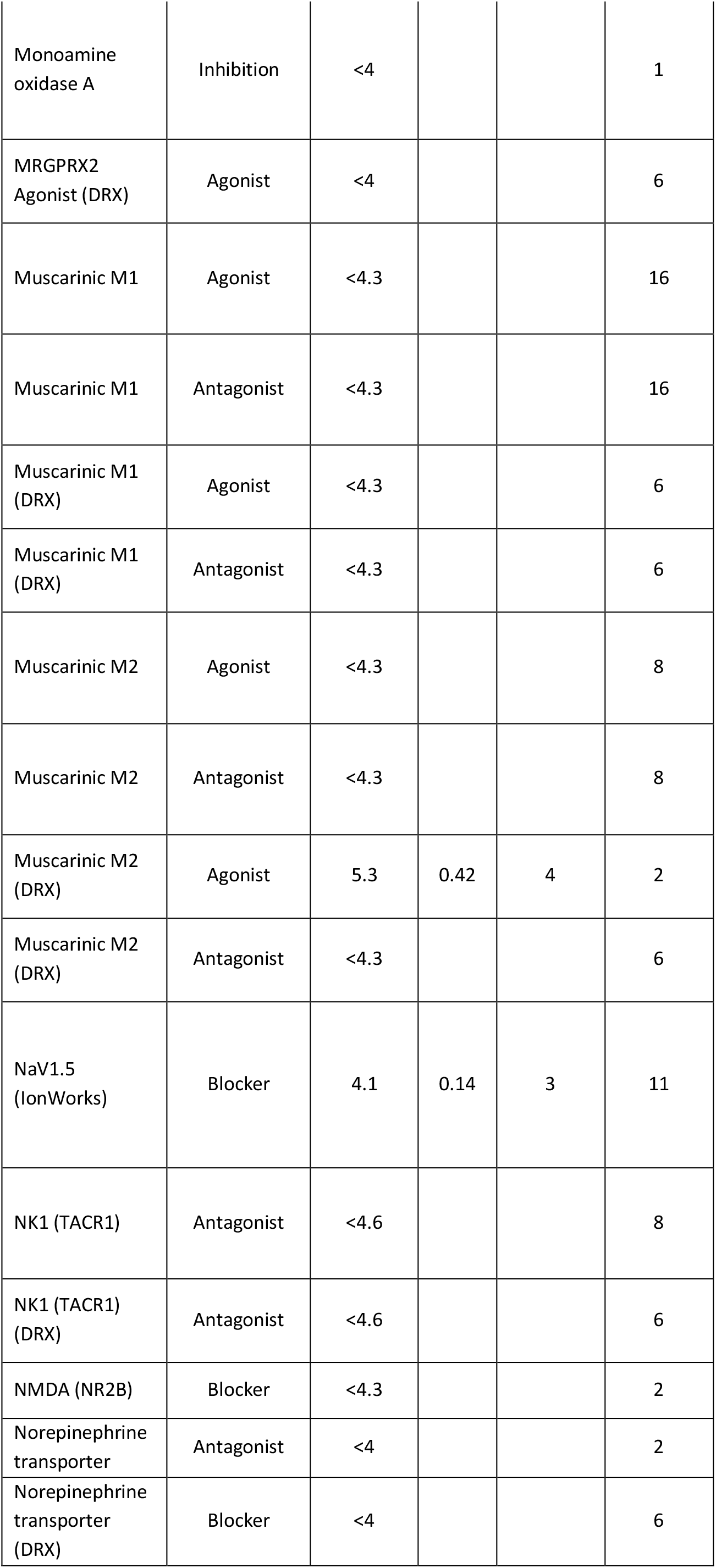
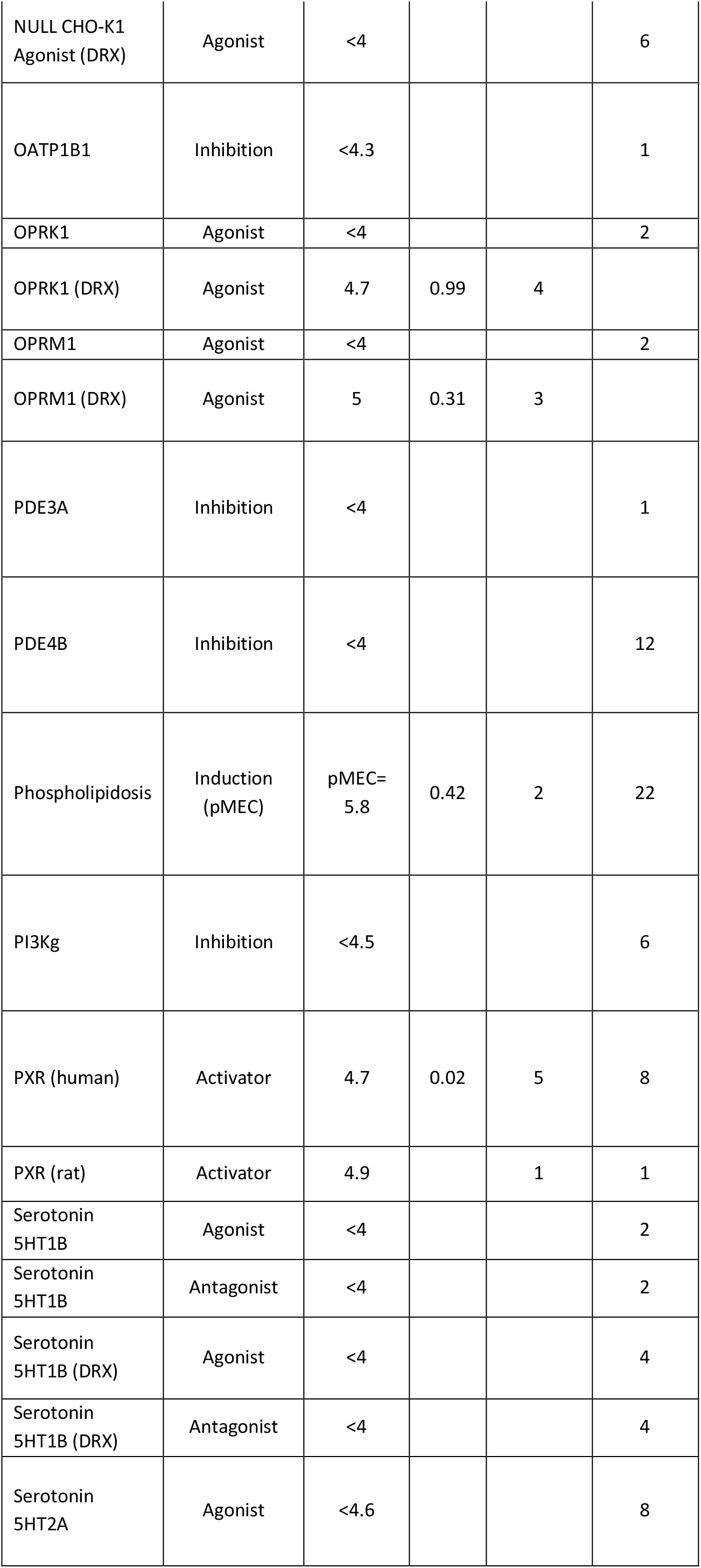
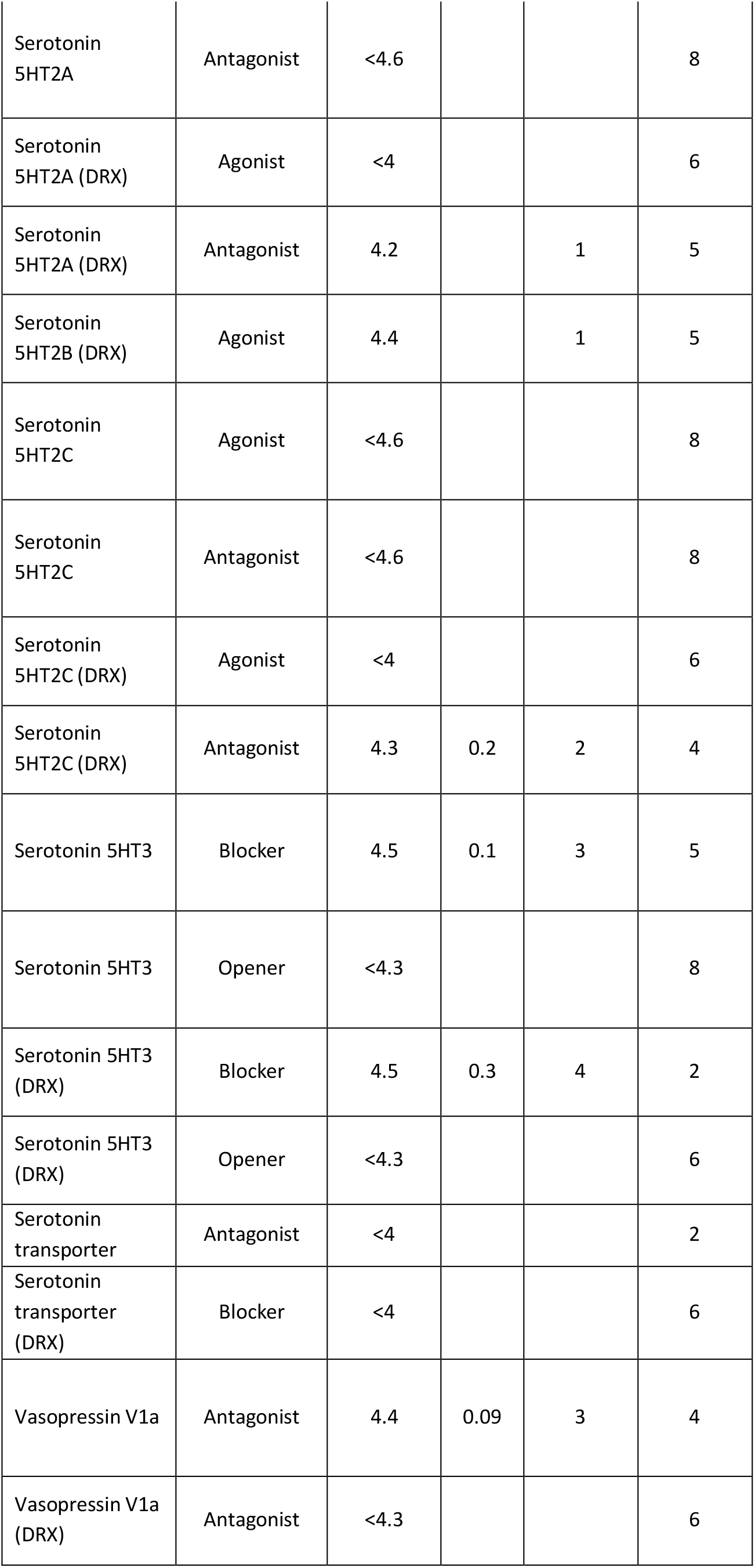

| ASSAY NAME | Mode | Mean (PXC50) | s.d. | Number of active assay replicates | Number of inactive assay replicates |
| --- | --- | --- | --- | --- | --- |
| α1 nicotinic AChR | Blocker | 4.5 | 0.14 | 3 | 1 |
| α1 nicotinic AChR | Opener | <4.3 | | | 8 |
| Acetyl-cholinesterase | Inhibition | 4.0 |  | 1 | 9 |
| Adenosine A2a | Agonist | <4 |  |  | 1 |
| Adrenergic α1B | Antagonist | <4.6 | | | 8 |
| Adrenergic α1B (DRX) | Antagonist | <4.6 | | | 6 |
| Adrenergic α2C | Agonist | <4 | | | 2 |
| Adrenergic α2C (DRX) | Agonist | <4 | | | 6 |
| Adrenergic β2 | Agonist | <4 | | | 2 |
| Adrenergic β2 | Antagonist | <4 | | | 2 |
| Adrenergic β2 (DRX) | Agonist | <4 | | | 6 |
| Adrenergic β2 (DRX) | Antagonist | <4 | | | 6 |
| Aryl hydrocarbon receptor | Activator | <4 |  |  | 1 |
| Aurora B (STK12) | Inhibition | <4.5 |  |  | 1 |
| Bile Salt Export Pump | Inhibition | 4 | 0.7 | 4 | 6 |
| Cannabinoid CB1 (DRX) | Agonist | <4 |  |  | 4 |
| Cannabinoid CB2 | Agonist | <4 |  |  | 8 |
| Cell Health | Membrane Permeability | <3.7 |  |  | 11 |
| Cell Health | Mitochondrial Potential | 3.8 | 0.3 | 6 | 5 |
| Cell Health | Nucleus (size) | <3.7 |  |  | 11 |
| Cyclooxygenase2 (COX2) | Inhibition | <4 |  |  | 1 |
| Dopamine D1 | Antagonist | <4 |  |  | 2 |
| Dopamine D1 (DRX) | Antagonist | <4 |  |  | 6 |
| Dopamine D2 | Agonist | <4 |  |  | 2 |
| Dopamine D2 | Antagonist | <4 |  |  | 2 |
| Dopamine D2 (DRD2S) (DRX) | Agonist | <4 |  |  | 6 |
| Dopamine D2 (DRD2S) (DRX) | Antagonist | <4 |  |  | 6 |
| GABA-A (Barracuda) | Agonist | <4 |  |  | 25 |
| GABA-A<br>(Barracuda) | Antagonist | <4 |  |  | 16 |
| GABA-A<br>(Barracuda) | Pos Mod | <4 |  |  | 13 |
| Glucocorticoid<br>Receptor | Agonist | <4 |  |  | 1 |
| hERG<br>(Barracuda) | Blocker | 4.8 |  | 1 | 19 |
| Histamine H1 | Antagonist | <4.6 |  |  | 8 |
| Histamine H1<br>(DRX) | Antagonist | 5 | 0.9 | 2 | 4 |
| KCNQ1/minK | Blocker | <4.6 |  |  | 12 |
| Kv1.5 | Blocker | <4.3 |  |  | 12 |
| L-type Ca<br>channel<br>(CaV1.2) | Blocker | >5.67 |  | 2 | 10 |
| LCK | Inhibition | <4.5 |  |  | 1 |
| Monoamine oxidase A | Inhibition | <4 |  |  | 1 |
| MRGPRX2 Agonist (DRX) | Agonist | <4 |  |  | 6 |
| Muscarinic M1 | Agonist | <4.3 |  |  | 16 |
| Muscarinic M1 | Antagonist | <4.3 |  |  | 16 |
| Muscarinic M1 (DRX) | Agonist | <4.3 |  |  | 6 |
| Muscarinic M1 (DRX) | Antagonist | <4.3 |  |  | 6 |
| Muscarinic M2 | Agonist | <4.3 |  |  | 8 |
| Muscarinic M2 | Antagonist | <4.3 |  |  | 8 |
| Muscarinic M2 (DRX) | Agonist | 5.3 | 0.42 | 4 | 2 |
| Muscarinic M2 (DRX) | Antagonist | <4.3 |  |  | 6 |
| NaV1.5 (IonWorks) | Blocker | 4.1 | 0.14 | 3 | 11 |
| NK1 (TACR1) | Antagonist | <4.6 |  |  | 8 |
| NK1 (TACR1) (DRX) | Antagonist | <4.6 |  |  | 6 |
| NMDA (NR2B) | Blocker | <4.3 |  |  | 2 |
| Norepinephrine transporter | Antagonist | <4 |  |  | 2 |
| Norepinephrine transporter (DRX) | Blocker | <4 |  |  | 6 |
| NULL CHO-K1 Agonist (DRX) | Agonist | <4 |  |  | 6 |
| OATP1B1 | Inhibition | <4.3 |  |  | 1 |
| OPRK1 | Agonist | <4 |  |  | 2 |
| OPRK1 (DRX) | Agonist | 4.7 | 0.99 | 4 |  |
| OPRM1 | Agonist | <4 |  |  | 2 |
| OPRM1 (DRX) | Agonist | 5 | 0.31 | 3 |  |
| PDE3A | Inhibition | <4 |  |  | 1 |
| PDE4B | Inhibition | <4 |  |  | 12 |
| Phospholipidosis | Induction (pMEC) | pMEC= 5.8 | 0.42 | 2 | 22 |
| PI3Kg | Inhibition | <4.5 |  |  | 6 |
| PXR (human) | Activator | 4.7 | 0.02 | 5 | 8 |
| PXR (rat) | Activator | 4.9 |  | 1 | 1 |
| Serotonin 5HT1B | Agonist | <4 |  |  | 2 |
| Serotonin 5HT1B | Antagonist | <4 |  |  | 2 |
| Serotonin 5HT1B (DRX) | Agonist | <4 |  |  | 4 |
| Serotonin 5HT1B (DRX) | Antagonist | <4 |  |  | 4 |
| Serotonin 5HT2A | Agonist | <4.6 |  |  | 8 |
| Serotonin 5HT2A | Antagonist | <4.6 |  |  | 8 |
| Serotonin 5HT2A (DRX) | Agonist | <4 |  |  | 6 |
| Serotonin 5HT2A (DRX) | Antagonist | 4.2 |  | 1 | 5 |
| Serotonin 5HT2B (DRX) | Agonist | 4.4 |  | 1 | 5 |
| Serotonin 5HT2C | Agonist | <4.6 |  |  | 8 |
| Serotonin 5HT2C | Antagonist | <4.6 |  |  | 8 |
| Serotonin 5HT2C (DRX) | Agonist | <4 |  |  | 6 |
| Serotonin 5HT2C (DRX) | Antagonist | 4.3 | 0.2 | 2 | 4 |
| Serotonin 5HT3 | Blocker | 4.5 | 0.1 | 3 | 5 |
| Serotonin 5HT3 | Opener | <4.3 |  |  | 8 |
| Serotonin 5HT3 (DRX) | Blocker | 4.5 | 0.3 | 4 | 2 |
| Serotonin 5HT3 (DRX) | Opener | <4.3 |  |  | 6 |
| Serotonin transporter | Antagonist | <4 |  |  | 2 |
| Serotonin transporter (DRX) | Blocker | <4 |  |  | 6 |
| Vasopressin V1a | Antagonist | 4.4 | 0.09 | 3 | 4 |
| Vasopressin V1a (DRX) | Antagonist | <4.3 |  |  | 6 |

**Supplementary Figure 1:**
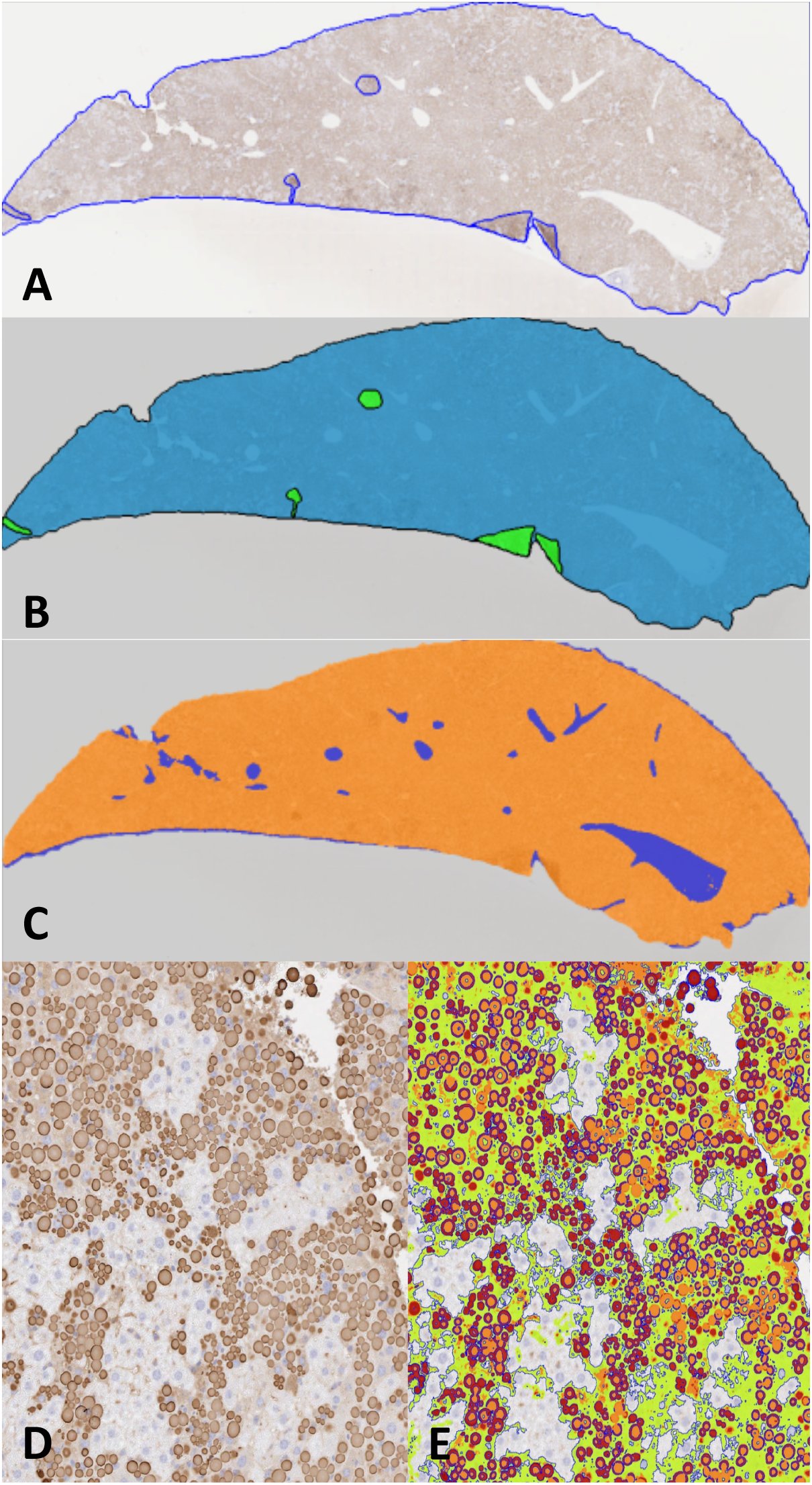
Image analysis. **A:** Liver section stained with 2C1 IHC with perimeter and artefacts drawn around freehand. **B:** Mask of A, green areas are excluded from quantification. **C**: Mask of A with white space (blue areas) removed from quantification. **D:** Liver section stained with 2C1 IHC. **E**: Section in D marked up using thresholds set by a Board-Certified pathologist. High intensity (red) and medium intensity (orange) areas describe the larger cytoplasmic inclusions and low intensity (green) areas describe more diffuse cytoplasmic staining (smaller polymers). Algorithm created using Definiens tissue studio version 2.7 software while blind to sample identity.

## References

1. D. A. Lomas, J. R. Hurst, B. Gooptu, Update on alpha-1 antitrypsin deficiency: new therapies. J Hepatol 65, 413–424. (2016).

2. Y. Wu, M. T. Swulius, K. W. Moremen, R. N. Sifers, Elucidation of the molecular logic by which misfolded α1-antitrypsin is preferentially selected for degradation. Proc. Natl. Acad. Sci. USA 100, 8229–8234 (2003).

3. A. Le, G. A. Ferrell, D. S. Dishon, A. L. Quyen-Quyen, R. N. Sifers, Soluble aggregates of the human PiZ α1-antitrypsin variant are degraded within the endoplasmic reticulum by a mechanism sensitive to inhibitors of protein synthesis. J. Biol. Chem. 267, 1072–1080 (1992).

4. D. Qu, J. H. Teckman, S. Omura, D. H. Perlmutter, Degradation of a mutant secretory protein, α1-antitrypsin Z, in the endoplasmic reticulum requires proteosome activity. J. Biol. Chem. 271, 22791–22795 (1996).

5. J. H. Teckman et al., The proteasome participates in degradation of mutant α1-antitrypsin Z in the endoplasmic reticulum of hepatoma-derived hepatocytes. J. Biol. Chem. 276, 44865–44872 (2001).

6. J. H. Teckman, J. K. An, K. Blomenkamp, B. Schmidt, D. Perlmutter, Mitochondrial autophagy and injury in the liver in α1-antitrypsin deficiency. Am J Physiol Gastrointest Liver Physiol 286, G851–G862 (2004).

7. D. A. Lomas, D. L. Evans, J. T. Finch, R. W. Carrell, The mechanism of Z α1-antitrypsin accumulation in the liver. Nature 357, 605–607 (1992).

8. A. Ordóñez et al., Endoplasmic reticulum polymers impair luminal protein mobility and sensitize to cellular stress in alphα_1_-antitrypsin deficiency. Hepatology 57, 2049–2060 (2013).

9. P. Strnad et al., Heterozygous carriage of the alphα_1_-antitrypsin Pi*Z variant increases the risk to develop liver cirrhosis. Gut 68, 1099–1107 (2019).

10. T. R. Dafforn, R. Mahadeva, P. R. Elliott, P. Sivasothy, D. A. Lomas, A kinetic mechanism for the polymerisation of α1-antitrypsin. J. Biol. Chem. 274, 9548–9555 (1999).

11. B. Gooptu et al., Inactive conformation of the serpin α1-antichymotrypsin indicates two stage insertion of the reactive loop; implications for inhibitory function and conformational disease. Proc. Natl. Acad. Sci (USA) 97, 67–72 (2000).

12. M. P. Nyon et al., Structural dynamics associated with intermediate formation in an archetypal conformational disease. Structure 20, 504–512 (2012).

13. U. I. Ekeowa et al., Defining the mechanism of polymerization in the serpinopathies. Proc. Natl. Acad. Sci. USA 107, 17146–17151 (2010).

14. M. Yamasaki, T. J. Sendall, M. C. Pearce, J. C. Whisstock, J. A. Huntington, Molecular basis of α1-antitrypsin deficiency revealed by the structure of a domain-swapped trimer. EMBO Rep. 12, 1011–1017 (2011).

15. A. S. Knaupp, V. Levina, A. L. Robertson, M. C. Pearce, S. P. Bottomley, Kinetic instability of the serpin Z α1-antitrypsin promotes aggregation. J Mol Biol. 396, 375–383 (2010).

16. A. Ordóñez et al., A single-chain variable fragment intrabody prevents intracellular polymerisation of Z α1-antitrypsin. FASEB J. 29, 2667–2678 (2015).

17. N. Motamedi-Shad et al., An antibody that prevents serpin polymerisation acts by inducing a novel allosteric behaviour. Biochem J. 473, 3269–3290 (2016).

18. R. A. Goodnow Jr, C. E. Dumelin, A. D. Keefe, DNA-encoded chemistry: enabling the deeper sampling of chemical space. Nature Reviews Drug Discovery 16, 131–147 (2017).

19. D. A. Lomas, D. L. Evans, S. R. Stone, W.-S. W. Chang, R. W. Carrell, Effect of the Z mutation on the physical and inhibitory properties of α1-antitrypsin. Biochemistry 32, 500–508 (1993).

20. J. A. Irving et al., An antibody raised against a pathogenic serpin variant induces mutant-like behaviour in the wild-type protein. Biochem J. 468, 99–108 (2015).

21. E. Miranda et al., A novel monoclonal antibody to characterise pathogenic polymers in liver disease associated with α1-antitrypsin deficiency. Hepatology 52, 1078–1088 (2010).

22. K. Yusa et al., Targeted gene correction of α1-antitrypsin deficiency in induced pluripotent stem cells. Nature 478, 391–394 (2011).

23. J. C. Whisstock, R. Skinner, R. W. Carrell, A. M. Lesk, Conformational changes in serpins: I. The native and cleaved conformations of alpha(1)-antitrypsin. J. Mol. Biol. 295, 651–665 (2000).

24. J. A. Irving, I. Haq, J. A. Dickens, S. V. Faull, D. A. Lomas, Altered native stability is the dominant basis for susceptibility of α1-antitrypsin mutants to polymerization. Biochem. J. 460, 103–115 (2014).

25. E. L. James, S. P. Bottomley, The mechanism of α1-antitrypsin polymerization probed by fluorescence spectroscopy. Arch. Biochem. Biophys. 356, 296–300 (1998).

26. D. B. Hood, J. A. Huntingdon, P. G. W. Gettins, α1-proteinase inhibitor variant T345R. Influence of P14 residue on substrate and inhibitory pathways. Biochemistry 33, 8538–8547 (1994).

27. L. Tan et al., Circulating polymers in α1-antitrypsin deficiency. Eur Respir J. 43, 1501–1504 (2014).

28. T. Hidvegi et al., An autophagy-enhancing drug promotes degradation of mutant alphα_1_-antitrypsin Z and reduces hepatic fibrosis. Science 329, 229–232 (2010).

29. S. Guo et al., Antisense oligonucleotide treatment ameliorates alpha-1 antitrypsin-related liver disease in mice. J Clin Invest. 124, 251–261 (2014).

30. B. Gooptu, D. A. Lomas, Conformational pathology of the serpins – themes, variations and therapeutic strategies. Annu. Rev. Biochem. 78, 147–176 (2009).

31. M. A. Clark et al., Design, synthesis and selection of DNA-encoded smallmolecule libraries. Nat Chem Biol. 5, 647–654 (2009).

32. C. C. Arico-Muendel, From haystack to needle: finding value with DNA encoded library technology at GSK. Med. Chem. Commun., 7, 1898–1909 (2016).

33. J. A. Irving et al., The serpinopathies: studying serpin polymerization in vivo. Methods Enzymol. 501, 421–466 (2011).

34. I. Haq et al., Reactive centre loop mutants of α1-antitrypsin reveal position-specific effects on intermediate formation along the polymerization pathway. Biosci Rep. 33, e00046 (2013).

35. I. Haq et al., Deficiency mutations of α1-antitrypsin differentially affect folding, function and polymerization. Am. J. Resp. Cell Mol. Biol. 2016, 71–80 (2016).

36. D. Kim, M.-H. Yu, Folding pathway of human α1-antitrypsin : characterisation of an intermediate that is active but prone to aggregation. Biochem. Biophys. Res. Commun. 226, 378–384 (1996).

37. J. E. Nettleship, J. Brown, M. R. Groves, A. Geerlof, Methods for protein characterization by mass spectrometry, thermal shift (ThermoFluor) assay, and multiangle or static light scattering. Methods Mol Biol. 426, 299–318 (2008).

38. A. J. McCoy et al., Phaser crystallographic software. J. Appl. Cryst. 40, 658–674 (2007).

39. P. Emsley, B. Lohkamp, W. G. Scott, K. Cowtan, Features and development of Coot. Acta Crystallogr D Biol Crystallogr. 66(Pt 4), 486–501 (2010).

40. G. N. Murshudov, A. A. Vagin, E. J. Dodson, Refinement of macromolecular structures by the maximum-likelihood method. Acta. Cryst. D53, 240–255 (1997).

41. K. Valkó, C. Bevan, D. Reynolds, Chromatographic Hydrophobicity Index by Fast-Gradient RP-HPLC: A high-throughput alternative to log P/log D. Anal. Chem. 69, 2022–2029 (1997).

42. M. C. Pearce et al., Preventing Serpin Aggregation: The molecular mechanism of citrate action upon antitrypsin unfolding. Protein Sci. 17, 2127–2133 (2008).

